# ATLAS: a package for multi-omic single-cell trajectory inference

**DOI:** 10.64898/2026.05.23.727175

**Authors:** Alessia Leclercq, Lorenzo Martini, Roberta Bardini, Alessandro Savino, Stefano Di Carlo

## Abstract

Single-cell trajectory inference is widely used to study cellular differentiation and fate decisions, yet most methods rely solely on transcriptomic data and therefore capture only part of the regulatory processes underlying cell-state transitions. Here we present ATLAS (Advanced Trajectory Learning from multi-omics At Single-cell resolution), a scverse-compatible Python package for trajectory inference from paired single-cell RNA-seq and ATAC-seq data. ATLAS integrates transcriptomic and chromatin accessibility information through Weighted Nearest Neighbor graphs, enabling both modalities to jointly inform pseudotime estimation, terminal-state identification, and fate probability inference within a unified multi-omic representation. Across synthetic and real datasets, ATLAS reconstructs coherent developmental trajectories, captures progressive fate commitment, and resolves biologically meaningful lineage structures, highlighting the value of multi-omic integration for characterizing cellular developmental dynamics. In addition, ATLAS enables joint analysis of transcription factor expression and accessibility-derived target-gene activity along pseudotime, providing insights into regulatory programs spanning transcriptomic and epigenomic layers that are not readily detectable from unimodal data. As a proof of concept, ATLAS recapitulates known hair follicle regulatory programs and reveals coherent multi-omic trajectories in which Lef1-associated regulatory patterns are linked to hair shaft differentiation. Overall, ATLAS provides an interoperable and biologically informative framework for studying cellular differentiation and regulatory dynamics in single-cell multi-omics experiments.

## Introduction

Trajectory Inference (TI) and pseudotemporal ordering have become central tools in single-cell transcriptomics for reconstructing dynamic biological processes such as differentiation, development, and cell fate decisions from snapshot data [6, 30, 31]. By ordering cells along putative developmental paths on the basis of their molecular profiles, rather than partitioning them into discrete clusters, TI provides a framework for studying continuous cellular processes and lineage relationships in complex systems [7].

Most existing TI methods were developed primarily for single-cell RNA sequencing (scRNA-seq) data. RNA-velocity approaches use spliced and unspliced transcript abundances to infer local cellular transitions [14, 3, 16], whereas pseudotime-based methods model progression along low-dimensional transcriptomic manifolds [30]. Widely used tools such as Palantir [24] and CellRank [15, 35] exemplify probabilistic and graph-based strategies for trajectory inference and fate prediction. However, these approaches rely almost exclusively on transcriptional information and therefore capture only one layer of cellular regulation.

Single-cell multiome technologies now enable simultaneous profiling of multiple molecular layers, mainly scRNA-seq and single-cell assays for transposase-accessible chromatin sequencing (scATAC-seq), at single-cell resolution [36, 12, 18]. These data provide complementary information on cell identity and regulatory programs, spanning both transcriptomic and epigenomic layers [28, 11]. Chromatin accessibility represents a permissive regulatory layer that can precede, accompany, or constrain transcriptional activation: although accessibility at a locus does not necessarily imply gene expression, changes in the accessibility of promoters and distal regulatory elements can mark genes that are competent for future activation. In this context, specific Transcription Factors (TFs), including Pioneer Transcription Factors (pTFs), can initiate or stabilize chromatin-accessibility changes at lineage-associated regulatory regions, thereby contributing to lineage priming and subsequent transcriptional commitment [20, 2]. Joint analysis of transcriptomic and chromatin-accessibility profiles can therefore reveal regulatory dynamics that would remain only partially resolved by either modality alone.

As a result, integrating RNA and ATAC modalities has become an active area of research [23]. Several alignment and latent-space integration frameworks have been proposed [26, 13], and among them the Weighted Nearest Neighbors (WNN) approach [10] has become widely adopted for multi-omic clustering, visualization, and cell type annotation [37].

Despite these advances, explicit integration of multi-omic information into trajectory inference remains limited. Most TI methods still estimate pseudotime and fate probabilities from RNA-derived representations, while chromatin accessibility is used mainly for auxiliary or post hoc interpretation. Extensions of RNA-velocity have incorporated paired RNA and ATAC measurements to model regulatory influences on transcriptional dynamics [16], but these methods remain fundamentally velocity-driven. They require more computationally expensive datasets and retain the main limitations of velocity modeling, including low signal-to-noise ratios and inconsistent results [4, 25]. Their applicability therefore remains restricted.

Alongside methodological advances, usability, interoperability, and community adoption have become critical for computational methods in single-cell analysis. The scverse ecosystem [33], built around shared data structures such as `AnnData` [34] and `MuData` [5], has emerged as a consortium of foundational tools for scalable and reproducible single-cell analysis.

Against this background, we present ATLAS, a scverse-compatible Python package for multi-omic trajectory inference from paired RNA and ATAC single-cell data. Rather than introducing a new trajectory-inference algorithm, ATLAS provides a standardized scverse-compatible framework that adapts established graph-based TI methods to WNN-based representations and couples multi-omic trajectory reconstruction with downstream regulatory visualization.

ATLAS extends established RNA-centric frameworks, including Palantir and CellRank, to operate on integrated multi-omic representations, allowing pseudotime ordering, terminal-state identification, and fate probability inference to be informed jointly by transcriptional and chromatin-accessibility-derived signals. ATLAS jointly analyses TF expression and target-gene activity along pseudotime, enabling the observation of coordinated changes across the two layers, including patterns consistent with TF-associated chromatin remodeling and lineage priming. By embedding multi-omic trajectory inference within the scverse ecosystem, ATLAS ensures interoperability with community standards and facilitates integration into existing workflows, enabling routine use of chromatin accessibility together with transcriptional data to study cellular differentiation dynamics in single-cell experiments.

## Methods

ATLAS is a Python package for trajectory inference in single-cell multiomics data that integrates gene expression, gene activity, and downstream trajectory analysis within a unified workflow (Fig. 1). Given paired scRNA-seq and scATAC-seq data organized in a multimodal representation, ATLAS reconstructs continuous developmental dynamics by extending existing graph-based TI methods [24, 35] within scverse. The framework also provides visualization tools for exploring inferred trajectories and multi-omic regulatory dynamics. The methodological components underlying these functionalities are described in the following sections.

**Fig. 1.**
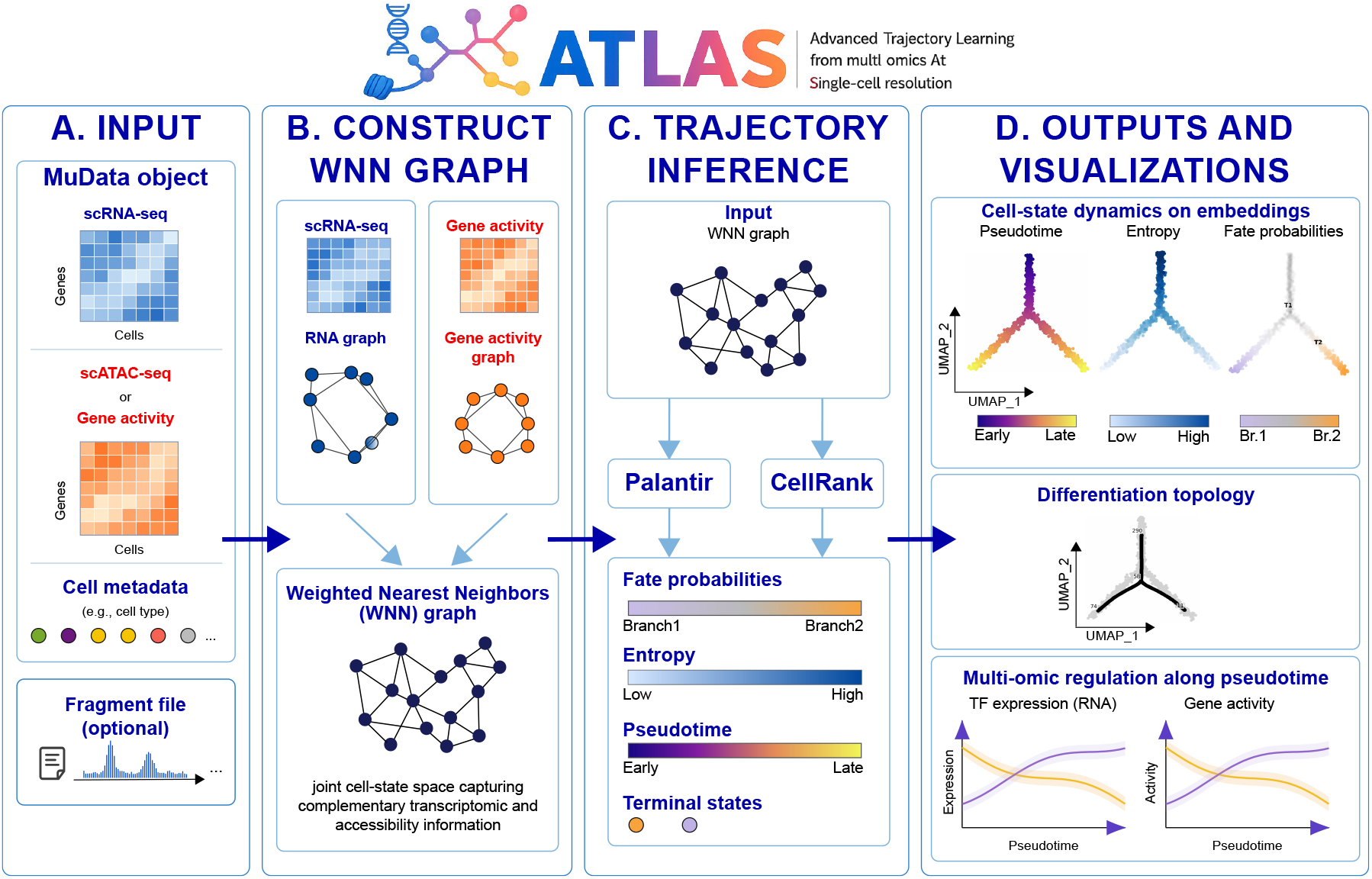
Overview of the Advanced Trajectory Learning from multi-omics At Single-cell resolution (ATLAS) workflow. (a) ATLAS takes as input paired scRNA-seq and scATAC-seq data, or a precomputed gene activity matrix, organized as a MuData object together with shared cell-level metadata. When scATAC-seq data are provided, a fragment file can be used to compute gene activity. (b) Gene expression and gene activity are integrated into a WNN graph, which defines a multi-omic cell-state representation. (c) The integrated representation is used for trajectory inference (d) ATLAS returns downstream visualizations for trajectory analysis and joint plots of TF expression and gene activity along pseudotime.

**Fig. 2.**
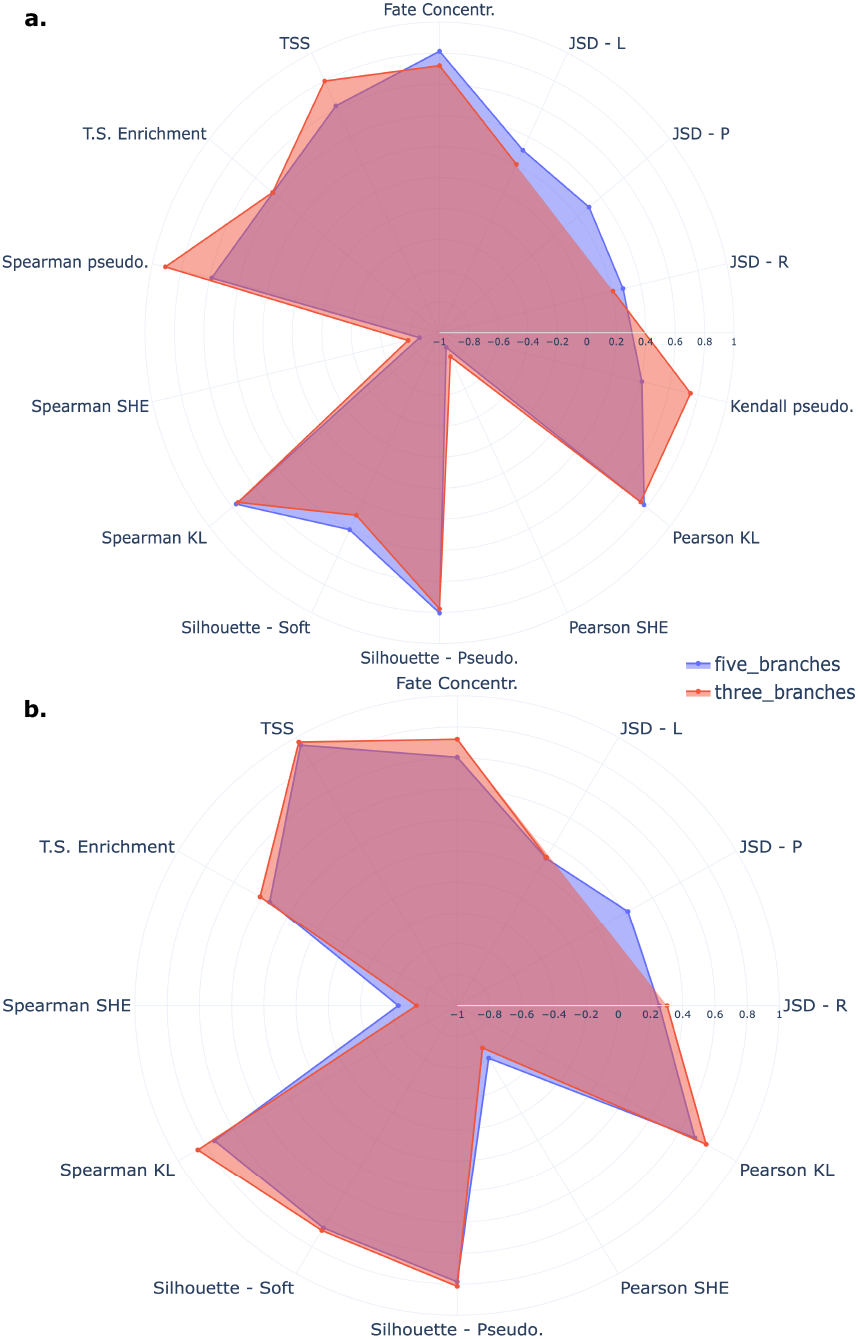
Median ATLAS performance on synthetic datasets. (a) Palantir-based trajectory inference. (b) CellRank-based trajectory inference.

### Workflow overview

Existing TI methods are typically designed for single-modality data and rely primarily on transcriptomic information. ATLAS addresses this limitation while preserving interoperability within the scverse ecosystem. Specifically, it operates on `MuData` objects [5], the scverse standard for paired single-cell multiomics data, which combine multiple single-modality structures [34] with shared cell-level metadata and cross-modality representations. This design enables integrated trajectory inference within existing single-cell analysis pipelines with minimal overhead (Supplementary Sec. 3).

The current implementation requires a `MuData` object containing a gene-expression modality and either a pre-computed gene activity matrix, where gene activity denotes a gene-level proxy of transcriptional potential derived from chromatin accessibility profiles [27, 21], or an scATAC-seq modality from which gene activity can be computed automatically using an auxiliary fragment file (Fig. 1a; Supplementary Sec. 1). As gene activity summarizes chromatin accessibility associated with genes rather than direct transcriptional output, it is interpreted throughout ATLAS as an accessibility-derived proxy of regulatory potential, and not as evidence of gene expression or causal TF-target regulation.

For trajectory inference, ATLAS builds on Palantir [24] and CellRank [15, 35], two graph-based methods that model cell-state transitions on neighborhood graphs. Because these frameworks operate on graph geometry rather than on a specific modality, they can be extended to integrated multiomic representations. ATLAS therefore replaces transcriptome-only neighborhood graphs with a WNN graph [10, 29] that combines gene expression and gene activity into a single representation (Fig. 1b; Supplementary Sec. 2). Compared with single-modality K-Nearest-Neighbors (KNN) graphs, WNN graphs adaptively weight each modality on a per-cell basis, allowing local neighborhoods to be defined by the most informative signal in each region of the manifold [10]. This strategy improves robustness to modality-specific noise and better preserves biologically meaningful local structure [29, 10].

After graph construction, ATLAS performs TI. In the Palantir-based workflow, the anisotropic kernel is adapted to the WNN graph before diffusion maps are computed on the multimodal manifold. In the CellRank-based workflow, the `PseudotimeKernel` object [35] orients the graph using pre-computed pseudotime values, after which Generalised Perron Cluster Cluster Analysis (GPCCA) [22] is applied to identify developmental branches.

ATLAS returns an updated `MuData` object containing pseudotime values, inferred terminal states, cell-fate probabilities, and entropy-based measures that summarize cellular plasticity and progressive fate commitment along pseudotime (Fig. 1c). Pseudotime is interpreted as an ordering of cells along an underlying developmental process rather than as an estimate of physical time [30, 14, 15, 24, 8]. Terminal states represent stable regions of the inferred differentiation dynamics toward which cells are probabilistically driven [24, 15, 35]. Cell-fate probabilities correspond to absorption probabilities in the inferred Markov process, and entropy-based measures summarize the uncertainty of fate commitment across terminal states.

To summarize the inferred developmental dynamics in a structured manner, scFates [8] is used to learn a principal graph representation of the process, providing a coherent topology that complements the probabilistic description given by cell-fate probabilities (Supplementary Fig. 1d,e).

Finally, to support interpretation of the inferred dynamics, ATLAS provides utilities to project cell-specific pseudotime, entropy measures, and fate probabilities onto low-dimensional embeddings (Fig. 1d). ATLAS also enables visualization of regulatory dynamics that are not accessible from transcriptomic data alone by jointly displaying TF expression and the activity profiles of putative target genes along pseudotime (Fig. 1d; Supplementary Sec. 4). Here, regulatory dynamics denote coordinated changes in TF expression and accessibility-derived target-gene activity along pseudotime, reflecting regulatory potential and transcriptional progression. Whereas gene expression captures transcriptional output, gene activity provides a proxy for regulatory engagement, enabling simultaneous interrogation of complementary regulatory layers. Lineage priming provides a pivotal example of the type of multi-omic regulatory pattern that ATLAS can help expose, as accessibility-derived gene activity may mark lineage-associated regulatory potential across developmental branches [20]. In this setting, regulatory programs involving pTFs are particularly relevant, as they link TF activity, chromatin accessibility, and cell-state competence [2].

### Datasets

Suitable datasets were selected to capture continuous biological processes with coherent progression from progenitor to terminal states and connected cellular neighborhoods along the differentiation continuum.

ATLAS was evaluated on both synthetic and real single-cell multi-omics datasets to assess performance under controlled conditions as well as in biologically complex settings.

Synthetic data were generated using scMultiSim [17], which simulates paired scRNA-seq and scATAC-seq profiles along predefined developmental trajectories and provides ground-truth pseudotime and lineage annotations (Suppl. Sec. 6).

Experiments on real data used three single-cell multi-omics datasets spanning distinct biological systems: SHARE-seq mouse hair follicle [20], Fresh Embryonic E18 Mouse Brain (5k) [9], and Human Fetal Brain [32] (Supplementary Sec. 8, 9, 10).

### Validation strategy

ATLAS was validated to assess whether the proposed multi-omic framework reconstructs coherent developmental trajectories. Comparisons against scRNA-seq-only approaches were also performed to evaluate the effect of multi-omic integration on trajectory inference results.

Validation is organized around two complementary objectives. The first evaluates trajectory inference accuracy when ground-truth information is available, using supervised metrics. The second assesses the confidence and internal consistency of inferred trajectories when explicit developmental ground truth is unavailable, using unsupervised metrics.

The supervised evaluation is performed on synthetic datasets, where the developmental structure is known. It assesses three main aspects of trajectory inference: pseudotime accuracy, terminal-state identification, and fate-dynamics reconstruction. Pseudotime accuracy is quantified through correlation-based agreement with ground-truth pseudotime values. Terminal-state inference is evaluated by measuring consistency with expected terminal populations and their position along the developmental process. Inferred cell-fate probabilities are then compared with expected transition probabilities derived from the predefined differentiation tree using the Jensen-Shannon divergence.

The unsupervised evaluation is applied in settings where explicit developmental ground truth is generally unavailable. In this case, validation focuses on the internal coherence and biological plausibility of the inferred trajectories. Three complementary aspects are assessed. First, correlation-based measures evaluate the relationship between cell plasticity and pseudotime, under the expectation that highly plastic cells occur at early developmental stages. Second, the evolution of cell-fate probability distributions along pseudotime is analyzed under the assumption that differentiation leads to progressive concentration toward terminal fates. Finally, silhouette metrics computed in the cell-fate probability space are used to assess the coherence and separation of inferred terminal states while accounting for the continuous nature of fate commitment.

Further details on the implementation of the evaluation metrics are provided in Supplementary Sec. 5 and Table 1. The benchmarking strategy against the scRNA-seq-only frameworks is described in Supplementary Sec. 11.

**Table 1.**
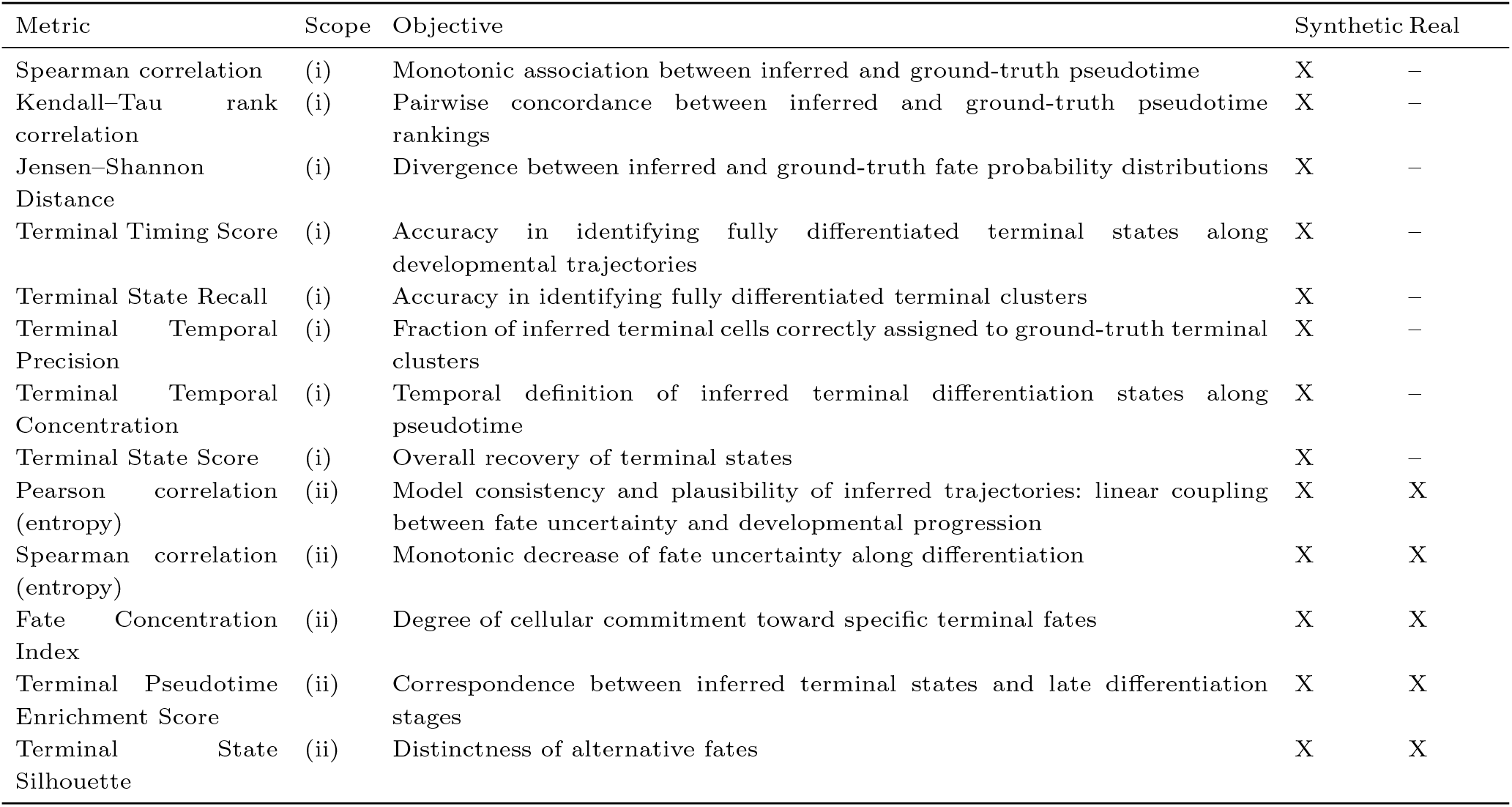
Overview of metric scope and applicability. Scope is either (i) a quantitative assessment of algorithm performance against ground truth, or (ii) an assessment of model consistency and plausibility of inferred trajectories without ground truth.

In real datasets, ATLAS was further examined for its ability to support cross-layer regulatory interpretation by visualizing branch-associated TF expression together with accessibility-derived target-gene activity. We used lineage priming and regulatory programs involving pTFs as a qualitative assessment of regulatory interpretability, because prior biological knowledge supports the presence of coordinated variation between TF activity, chromatin-accessibility-derived regulatory potential, and transcriptional progression along differentiation trajectories.

## Results and Discussion

### Synthetic Datasets

The synthetic benchmark comprised 25 paired scRNA-seq/scATAC-seq datasets of 1,000 cells each, generated across 3- and 5-branch topologies and multiple trajectory-ambiguity settings (Supplementary Sec. 6; Supplementary Table 1). Across all settings, ATLAS coherently recovered developmental structure at three complementary levels: pseudotime ordering, fate-probability dynamics, and terminal-state identification.

Pseudotime accuracy, assessed using Spearman’s *ρ* and Kendall’s *τ* against ground-truth pseudotime, was high in the 3-branch topology. In the 5-branch setting, median correlations decreased for both measures, with a stronger reduction in Kendall’s *τ* . This pattern suggests that higher branching complexity introduces more local ordering inconsistencies while largely preserving the global monotonic structure.

Fate-probability dynamics were also consistent with differentiation progression. Across simulations, pseudotime showed strong negative correlations with Shannon entropy and positive correlations with Kullback Leibler (KL) divergence and the Fate Concentration Index (FCI). These relationships were largely maintained in the 5-branch topology, with only mild attenuation in strength. Agreement with ground-truth fate distributions, measured by Jensen Shannon Divergence (JSD), remained overall good despite the cluster-level construction of the ground truth.

Terminal-state recovery was similarly coherent. The Terminal State Score indicated accurate localization of terminal cells, appropriate phenotypic purity, and substantial coverage of ground-truth terminal populations. Unsupervised metrics supported the same conclusion: the Terminal Pseudotime Enrichment Score was consistently positive, showing that inferred terminal states were enriched at late pseudotime values, and silhouette-based metrics indicated meaningful probabilistic separation of terminal outcomes across all settings. Increased branching complexity led to moderately higher dispersion in several terminal-state metrics.

Differences between the two trajectory-inference back ends emerged when variability across simulations was considered. The Palantir-based implementation showed slightly higher median performance, but also broader dispersion and occasional outliers. In contrast, the CellRank-based implementation generally showed lower variability across pseudotime, fate, and terminal-state metrics, despite marginally lower medians in some cases, suggesting greater robustness across simulations.

### Real Datasets

Results on the SHARE-seq mouse hair follicle dataset show that ATLAS disentangles the hair follicle developmental trajectory. As shown in Fig. 3c, ATLAS assigns a coherent pseudotemporal ordering to cells, recapitulating the known biological progression from Transient Amplifying Cells (TAC) toward Hair Shaft (HS) lineages (Medulla and Cuticle Cortex) and Inner Root Sheath (IRS) cells (Fig. 3b,c). Both trajectory-inference strategies identify the main developmental branches corresponding to the Cuticle Cortex and Medulla lineages (Fig. 3d,f; Supplementary Fig. 5d). When CellRank is used as the underlying trajectory-inference method, an additional terminal state associated with TAC cells is detected. Although this state is largely composed of cells with high pseudotime values, it does not correspond to a biologically expected endpoint (Fig. 3f; Supplementary Fig. 5d).

**Fig. 3.**
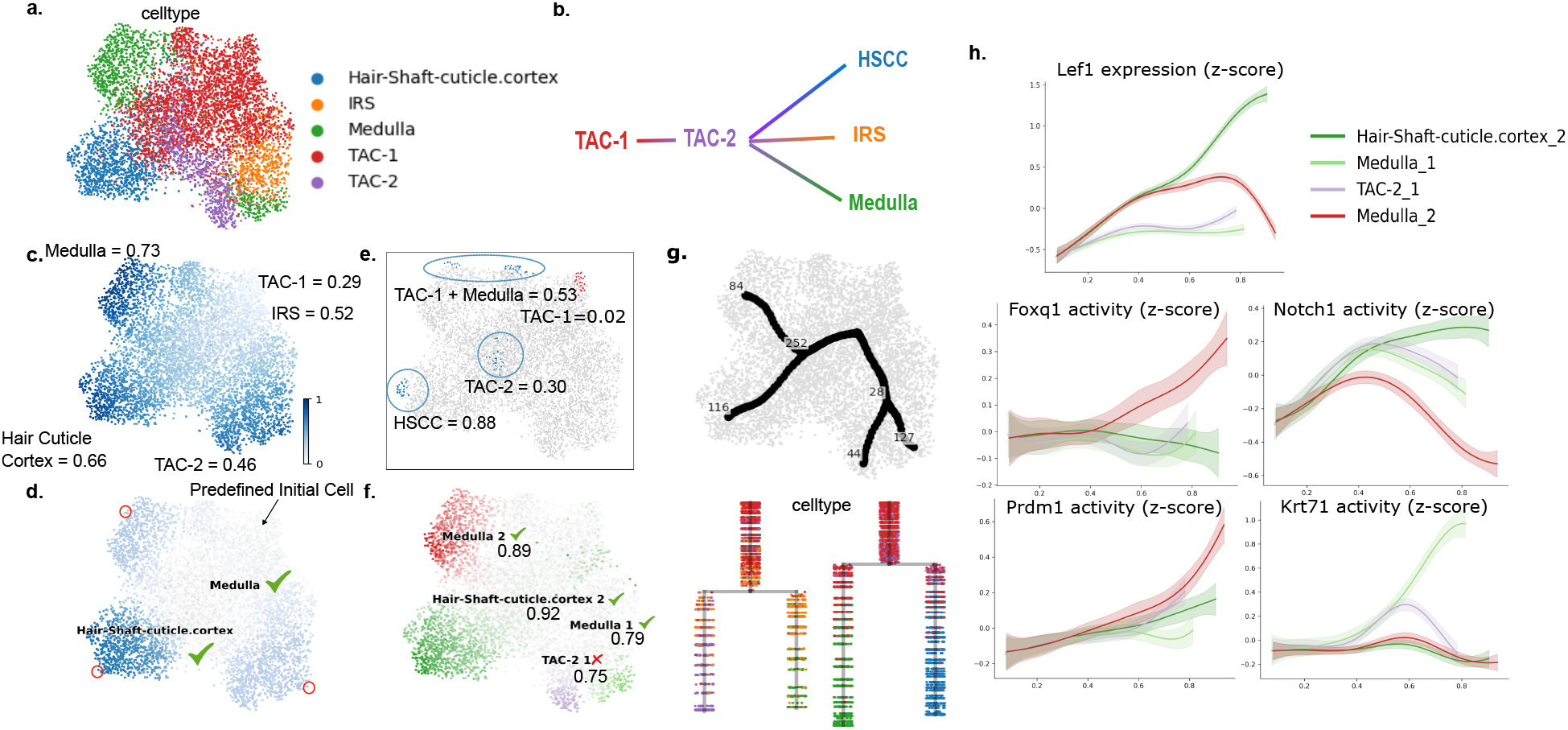
Overview of ATLAS results on the mouse skin dataset. (a) Cell-type annotations over the UMAP embedding. (b) Expected developmental trajectories. (c) Inferred pseudotime using ATLAS and average pseudotime for each cell cluster. (d) Terminal states (red) and fate probabilities inferred using the Palantir-based workflow. (e) Initial (red) and intermediate (blue) macrostates with average pseudotime. (f) Terminal macrostates and developmental branches inferred using the CellRank-based workflow. (g) Graph plots and dendrograms obtained by applying scFates to the fate-probability matrix in (f); colors correspond to cell-type annotations. (h) Lef1 expression and accessibility-derived target-gene activity (Foxq1, Prdm1, Notch1, Krt71) along the inferred developmental lineages.

Increasing the number of macrostates improves resolution in the identification of the initial population, which is assigned to early TAC cells (average pseudotime 0.02), and of multiple intermediate states corresponding to transitional cell types or populations with low average pseudotime (Fig. 3e). Notably, IRS cells are not inferred as a terminal state. Instead, the developmental dendrograms from both ATLAS-CellRank experiments (Fig. 3g; Supplementary Fig. 5f) indicate that IRS cells are transitional.

The consistency of these results is supported by unsupervised evaluation metrics (Supplementary Table 5). Across all experiments, pseudotime and entropy show strong concordance, with correlation magnitudes around 0.8. ATLAS also achieves positive Terminal Pseudotime Enrichment Score (TPES), indicating enrichment of inferred terminal states at late differentiation stages. In addition, fate probabilities progressively concentrate along pseudotime (FCI around 0.83– 0.85), reflecting increasing cellular commitment. Relatively low silhouette scores indicate gradual and overlapping fate transitions, consistent with the observed TPES and the progressive increase in FCI, and therefore support a continuous lineage-commitment process.

The SHARE-seq dataset also highlights the added value of ATLAS for disentangling epigenomic and transcriptomic regulatory dynamics along developmental branches. Joint visualization of TF expression and accessibility-derived target-gene activity reveals patterns that are not accessible from transcriptomic data alone. As a representative example, Lef1, a TF implicated in chromatin remodeling during hair follicle development [20, 1, 38], and a set of selected putative target genes were examined. The expression profile of Lef1 along the inferred developmental branches (Fig. 3h) recapitulates the expected increasing trend along HS-related lineages [20]. In the same branches, the activity profiles of Foxq1 and Prdm1 show increased accessibility-derived activity, consistent with a Lef1-associated regulatory program compatible with lineage priming during HS differentiation. In parallel, Notch1 displays decreasing activity along the inferred pseudotemporal ordering, in agreement with its known association with IRS-related regulatory programs [20].

Lef1-associated regulation of keratin genes [1] is also reflected in the observed patterns. Previous studies [20] reported increased chromatin accessibility at Krt71 in IRS cells. In Fig. 3h, Krt71 activity is enriched in the “Medulla-1” and “TAC-2 1” branches, suggesting that these trajectories capture IRS-like regulatory features. Altogether, these multiomic signals are consistent with established biological knowledge, further supporting the biological interpretability of ATLAS-inferred trajectories and highlighting its ability to uncover coherent regulatory programs through integrated multiomic .

On the Embryonic Mouse Brain dataset (Supplementary Sec. 8), pseudotime inference assigns the lowest pseudotime values to the cluster containing Radial Glia (RG) and Oligodendrocyte Precursor Cells (OPC) cells. ATLAS also orders clusters along the neuronal lineage by assigning increasing pseudotime values to the Intermediate Precursor Cells (IPC), Ventricular - Subventricular Zone (V-SVZ), and then the cortical layers (Subplate cells, Upper Layer neurons, and Deeper Layer neurons) [19]. Cluster-average pseudotime values further show that the expected terminal clusters are placed at similar developmental stages across lineages, with average pseudotime values ranging from 0.42 to 0.49 (Supplementary Fig. 6c).

The choice of trajectory-inference back end affects terminal-state recovery. The Palantir-based workflow recovers a single terminal state corresponding to Upper Layer neurons (Supplementary Fig. 6d). In contrast, the CellRank-based workflow steadily recovers terminal macrostates composed of both Upper and Deeper Layer neurons (Supplementary Figs. 6f,i), thereby resolving cortical layer development and neuronal specification into distinct layers [19]. As in the SHARE-seq dataset, increasing the number of macrostates improves the identification of the initial population composed of RG cells and reveals additional macrostates corresponding to initial cells, intermediate clusters, or terminal clusters with lower pseudotime values than the main terminal groups (Supplementary Fig. 6h).

Metric results for this dataset are reported in Supplementary Table 6. ATLAS maintains a substantial correlation between pseudotime and entropy, with absolute values ranging from 0.6 to 0.7. Positive TPES values indicate that inferred terminal states remain enriched at late pseudotime. Consistently with the SHARE-seq results, silhouette metrics together with TPES and FCI support gradual fate commitment.

In experiments on the Human Fetal Brain dataset (Supplementary Sec. 10), ATLAS identifies terminal states corresponding to Subplate, Upper Layer, and Deeper Layer neurons, as well as a terminal state associated with the “RG, Astro” and “mGPC OPC” clusters. In contrast to the Embryonic Mouse Brain dataset, recovery of these terminal lineages is supported by the expression of canonical marker genes for OPC and astrocytes (Supplementary Figs. 9f–h), suggesting the presence of fully differentiated supporting cells in the dataset. In this case, the WNN graph contains a disconnected component composed of multipotent Glial Progenitor Cells (mGPC) cells that do not express canonical markers for either mGPC or OPC populations, as shown in Supplementary Figs. 9 h,j. Disconnected components are a known challenge for random walk–based algorithms because the behavior of the Markov chain depends on the connected component selected as the starting point.

This issue affects multiple components of ATLAS, including pseudotime inference (Supplementary Fig. 10c), macrostate identification, and developmental probabilities (Supplementary Fig. 10). Cells in the disconnected component are assigned the highest pseudotime values and are identified as a terminal lineage. Moreover, the disconnected structure likely induces a singleton in the GPCCA decomposition underlying CellRank, causing the “mGPC, OPC 1” macrostate to be selected as the initial state.

Removal of the disconnected component improves pseudotime inference, with Astrocytes, Oligodendrocytes (OG), Deeper Layer Neurons, and Subplate Neurons exhibiting the highest pseudotime values (Supplementary Fig. 12c). Furthermore, when Palantir is used as the underlying TI method, four of the five expected terminal states are identified (Supplementary Fig. 12d). The CellRank-based workflow also recovers the expected terminal macrostates and developmental lineages; however, increasing the number of macrostates to 10 yields a more fragmented representation of cortical layers (Supplementary Fig. 12i). This fragmentation is likely driven by the presence of multiple samples and substantial batch effects (Supplementary Fig. 9b), rather than by an intrinsic tendency toward over-segmentation.

Cycling progenitor cells, which are expected to represent the initial state of the differentiation process [16], are not explicitly identified as an initial macrostate. Nevertheless, the inferred trajectories remain consistent with the pseudotime ordering.

### Evaluation Against Original RNA-only Methods

ATLAS shows positive average shifts in rank-based pseudotime correlations and Terminal State Score as compared to the original scRNA-seq based TI methods across synthetic datasets and 3- and 5-branch topologies. Correlations between pseudotime and entropy are comparable between ATLAS and scRNA-seq–only methods (Δ_*median*_ between 0.01 and 0.13), and differences in FCI are generally modest, suggesting similar overall levels of fate concentration.

When terminal states are fixed, ATLAS achieves slightly better JSD, especially in the root cluster.

Overall, the paired evaluation on synthetic datasets indicates that ATLAS matches or surpasses the performance of established scRNA-seq–only TI methods across all evaluated aspects, with its most evident advantages emerging in more complex topologies where accurate trajectory reconstruction is particularly challenging. Supplementary Tables 3 and 4 summarize the performance differences between ATLAS and the original RNA-only methods across synthetic datasets (Supplementary Sec. 11).

In the SHARE-seq dataset, scRNA-seq–only TI recovers a global developmental ordering broadly comparable to ATLAS, and both Palantir and CellRank identify the major HS lineages (Supplementary Fig. 5). However, increasing the number of macrostates leads to progressive over-fragmentation rather than improved resolution of intermediate states. Several terminal macrostates become associated with TAC populations, including cells at low pseudotime values, and initial states are inconsistently assigned to differentiated populations such as Hair Shaft Cuticle Cortex (HSCC), Medulla, and IRS. Quantitative metrics (Supplementary Table 5) support this interpretation: although scRNA-seq–only methods show comparable entropy correlations and higher silhouette scores, TPES indicates frequent assignment of terminal states to early or intermediate pseudotime regions. This pattern suggests that the increased separation is driven by fragmentation rather than by biologically coherent lineage resolution.

Similar limitations emerge in the Embryonic Mouse Brain dataset. scRNA-seq–only CellRank analyses fragment fate probabilities across multiple terminal macrostates (Supplementary Fig. 7), with RG populations simultaneously identified as initial and terminal states and Neuronal Intermediate Progenitor Cells (nIPC) cells consistently recovered as terminal lineages. Increasing the number of macrostates further amplifies these inconsistencies, producing initial states with higher average pseudotime than intermediate populations. Palantir, in contrast, recovers only a single terminal lineage within the Deeper Layer cluster. Unsupervised metrics (Supplementary Table 6) support these observations: while entropy correlations are comparable or stronger in scRNA-seq–only analyses, only ATLAS consistently preserves coherent relationships between pseudotime, KL divergence, and TPES, indicating a more biologically consistent temporal organization of terminal states. In the Human Fetal Brain dataset, graph disconnection also affects scRNA-seq–only experiments (Supplementary Fig. 11), producing inference artifacts. In particular, Palantir fails to recover developmental lineages when relying solely on gene-expression data.

After removal of the disconnected component, several inconsistencies remain in scRNA-seq–only analyses (Supplementary Fig. 13). Low pseudotime values are assigned to Upper Layer neurons, while CellRank systematically identifies a terminal lineage within the IPC cluster, inconsistent with the biological role of this population. Increasing the number of macrostates again leads to over-fragmentation rather than improved identification of intermediate states, including multiple terminal macrostates associated with “mGPC, OPC” cells. Quantitative metrics evaluation (Supplementary Table 8) indicates overall higher confidence and consistency for the ATLAS-based TI strategy as compared to original RNA-only methods.

## Conclusion

This work presents ATLAS, a scverse-compatible Python package for trajectory inference that integrates transcriptomic and chromatin accessibility information through WNN-based representations.

Across synthetic and real datasets, ATLAS reconstructs coherent developmental trajectories while capturing progressive fate commitment and terminal differentiation. In particular, integrating chromatin accessibility improves the consistency of terminal-state identification in settings where scRNA-seq–only approaches produce fragmented or biologically inconsistent solutions. Challenges such as graph disconnection and batch effects remain relevant, but multi-omic integration mitigates their impact and yields a more informative representation of the cellular state space and differentiation dynamics.

Beyond trajectory reconstruction, ATLAS supports the exploration of regulatory dynamics by jointly visualizing TF expression and accessibility-derived target-gene activity along pseudotime. This enables the observation of branch-associated multi-omic patterns consistent with regulatory priming, including programs involving pTFs, without implying direct causal inference of TF-target relationships. In the SHARE-seq hair follicle dataset, this functionality recapitulated Lef1-associated regulatory patterns linked to hair shaft differentiation, illustrating how ATLAS can connect inferred developmental trajectories with biologically interpretable regulatory programs. Overall, ATLAS shows that integrating chromatin accessibility into trajectory inference improves robustness, interpretability, and biological consistency, supporting the use of multi-modal approaches to study dynamic processes in single-cell biology.

## Data and Code Availability Statement

Fresh Embryonic E18 Mouse Brain (5K) can be accessed at the 10x website at 10xgenomics.com. SHARE-seq mouse hair follicle and Human Fetal Brain datasets can be found at NCBI GEO, respectively under accession codes GSE140203 and GSE162170. Source code for ATLAS package is available on GitHub at https://github.com/smilies-polito/atlas-smilies, while code for experiments, figures and table generation can be found on GitHub at https://github.com/smilies-polito/atlas-experiments.

## Author contributions statement

We follow here the Contributor Roles Taxonomy (CRediT). AL: conceptualization, methodology, software, visualization, formal analysis, investigation, writing – original draft; LM: conceptualization, methodology, validation, investigation, writing – original draft, supervision; RB: conceptualization, methodology, supervision, validation, writing - review & editing, project administration; AS: supervision, writing - review & editing; SDC: supervision, writing - review & editing, project administration.

