## Supplementary Fig. for "ATLAS: a package for multi-omic single-cell trajectory inference"

#### 1 Gene Activity Computation

Advanced Trajectory Learning from multi-omics At Single-cell resolution (ATLAS) uses accessibility-derived gene activity instead of peak-level accessibility because gene-level aggregation better preserves the smooth neighborhood structure required by diffusion- and graph-based trajectory inference, whereas peak-level accessibility introduces local discontinuities that fragment the graph [1]. It computes gene activity scores with the `muon.atac.tl.count_fragments_features` function from muon [2] and then normalizes them to equalize total counts across cells.

Gene activity is therefore interpreted as an accessibility-derived proxy of regulatory potential associated with genes, rather than as direct transcriptional output or evidence of causal TF-target regulation.

This computation requires the single-cell assays for transposase-accessible chromatin sequencing (scATAC-seq) data and two additional inputs: a fragment file describing the mapping of DNA fragments to the reference genome, formatted according to muon requirements [2], and a feature dataframe defining gene-associated regions. At minimum, the feature dataframe must contain the columns `chromosome`, `start`, and `end`; users can optionally include a `strand` column to encode gene orientation.

#### 2 Building the Integrated Multi-omic Representation using ATLAS

ATLAS requires preprocessed single-cell RNA sequencing (scRNA-seq) data. In the default workflow, users log-normalize the expression matrix after standard quality-control filtering [3, 4]. ATLAS constructs unimodal K-Nearest Neighbors (KNN) graphs on reduced representations with the `scanpy.pp.neighbors` function from Scanpy [5]. By default, it applies Principal Component Analysis (PCA) to both scRNA-seq and gene-activity matrices, although users can provide alternative low-dimensional embeddings supported by Scanpy. It then computes the integrated Weighted Nearest Neighbors (WNN) graph with the `muon.pp.neighbors` function from muon.

The preprocessing step returns a separate `MuData` object containing the gene-expression and gene-activity modalities together with the metadata and multimodal structures inherited from the original object.

#### 3 Further Details on ATLAS TI

ATLAS implements multimodal trajectory inference through two wrapper classes, `PalantirExtension` and `CellRankExtension`. These classes expose the main parameters of the original methods through a unified scverse-compatible interface while reusing the corresponding package implementations.

Palantir- and CellRank-specific mathematical details remain unchanged and are described in the original publications [6, 7, 8, 9].

In addition, ATLAS computes two entropy-based quantities from the inferred cell-fate probabilities: Shannon entropy $H_c$ and Kullback Leibler (KL) divergence $KL_c$.

$$H_c := -\sum_{t \in \mathcal{T}} F_{c,t} * log(F_{c,t}) \tag{1}$$

$$KL_c := \sum_{t \in \mathcal{T}} F_{c,t} * log\left(\frac{F_{c,t}}{\bar{F}_t}\right) \quad (2)$$

Given a set of terminal states $\mathcal{T}$, Shannon entropy quantifies the uncertainty of the fate probability distribution for each cell $c$. Higher values indicate that fate probability mass is distributed across multiple terminal states, whereas lower values indicate preferential commitment toward a specific lineage.

KL divergence quantifies the divergence between the cell-specific fate probability distribution and a reference distribution defined by the population-wide average fate probabilities $\bar{F}$. While Shannon entropy measures the spread of probabilities within a cell, KL divergence measures how different that distribution is from the population average.

### 4 Downstream Visualisations and Analyses

ATLAS generates embedding-based visualizations of cell-wise pseudotime, entropy values, and most likely developmental lineages with ScVelo [10] (Supplementary Figure 1 b,c). These visualizations allow users to inspect the inferred developmental ordering, the progressive resolution of fate uncertainty, and the spatial organization of fate probabilities on low-dimensional embeddings.

To visualize branch-resolved regulatory dynamics, ATLAS provides utilities to jointly display Transcription Factor (TF) expression and accessibility-derived activity of putative target genes along pseudotime (Supplementary Fig. 1f). These visualizations are intended to support qualitative cross-layer interpretation of regulatory patterns, rather than to infer causal TF-target interactions. In particular, programs involving Pioneer Transcription Factors (pTFs) can be inspected as a representative case in which TF expression and chromatin-accessibility-derived regulatory potential are expected to vary coherently across developmental branches.

To visualize branch-resolved regulatory dynamics, ATLAS fits independent branch-specific models that account for soft cell-to-branch assignments through fate probabilities (Supplementary Figure 1f). Let $\tau_i \in [0, 1]$ denote the pseudotime of cell $i$, let $\mathcal{T}$ denote the set of inferred terminal states, and let $F_{i,t}$ denote the probability that cell $i$ differentiates along branch $t \in \mathcal{T}$. Let $y_i^{\mathrm{TF}}$ denote the expression of the TF and $y_i^G$ the activity of the downstream target gene in cell $i$. For each branch $t \in \mathcal{T}$ and for each biological signal $s$, ATLAS fits a Generalized Additive Model (GAM) of the form shown in Equation 3, where $f_t(\cdot)$ is a smooth branch-specific function of pseudotime represented with penalized regression splines.

$$\begin{aligned} \tilde{y}_i^s &= f_t^s(\tau_i) + \epsilon_i \\ \sum_i F_{i,t}(y_i^s - f_t^s(\tau_i))^2 &+ \lambda \int (f_t^{s\prime\prime}(\tau))^2 d\tau \end{aligned} \quad (3)$$

The first term measures weighted goodness-of-fit, whereas the second penalizes excessive curvature. The smoothing parameter $\lambda$ controls the trade-off between data fidelity and smoothness. The resulting curves provide branch-specific summaries of coordinated variation between TF expression and accessibility-derived gene activity along the inferred pseudotemporal ordering.

ATLAS also provides utilities for principal-tree and dendrogram inference through a wrapper around scFates [11] (Supplementary Figure 1 d,e). It uses the original scFates implementation without modification and supplies inferred fate probabilities to guide topology construction and branch assignment. Additional methodological details are available in the original publication [11].

### 5 Evaluation metrics

ATLAS defines metrics used to evaluate inferred pseudotime and developmental trajectories on simulated and real single-cell datasets. Simulated data provide ground-truth labels and therefore support supervised evaluation. Most real datasets do not provide developmental ground truth, so we additionally use unsupervised metrics to quantify the consistency and structure of the inferred landscape directly from the data.

In the following equations, $L = \{L_1, \ldots, L_d\}$ denotes the set of ground-truth terminal (*leaf*) clusters, $\tau$ denotes pseudotime, and $\mathcal{T} = \{\mathcal{T}_1, \ldots, \mathcal{T}_n\}$ denotes the set of inferred terminal states. The set $T = \bigcup_j \mathcal{T}_j$ contains all cells assigned to terminal states, and $m(\cdot)$ maps each cell to its ground-truth cluster. The fate-probability matrix is denoted by $F \in \mathbb{R}^{N \times K}$, where $F_{c,k}$ is the probability of cell $c$ committing to terminal state $k$. The parameter $\alpha \geq 0$ controls pseudotime weighting, $d(\cdot, \cdot)$ denotes distances in fate-probability space, and $w_{\cdot,\cdot}$ denotes the probability that two cells share the same terminal fate.

The supervised evaluation first measures agreement between inferred and ground-truth pseudotime using Spearman and Kendall-tau correlations. Spearman correlation evaluates monotonic agreement, whereas Kendall-tau measures concordance of pairwise cell orderings.

Agreement between inferred and ground-truth fate probability distributions is evaluated with the Jensen-Shannon Distance (JSD), defined as the square root of the Jensen-Shannon divergence computed with natural logarithms. To ensure comparability across methods that may infer different numbers or definitions of terminal states, terminal states are fixed to a predefined set of cells shared across all methods before fate probability estimation. Because the ground-truth fate probabilities defined in Supplementary Section 6 imply different interpretations of the JSD across developmental stages, evaluation is stratified by cluster type. Clusters are categorized as *root*, *parent*, or *leaf* (Supplementary Fig. 2 a), and the JSD is computed separately for each category.

Finally, supervised terminal-state metrics evaluate complementary aspects of terminal-state identification, including temporal placement, cluster composition, recovery of expected terminal populations, and temporal concentration.

- Terminal Timing Score (TTS): evaluates whether inferred terminal states are positioned at the end of the developmental trajectory (Equation 4). The final TTS score is the average of $TTS_i$ across ground-truth terminal clusters. Values lie in $[0, 1]$, and larger values indicate more accurate recovery of terminal timing.

$$
\begin{aligned}
&\mathcal{C}_i = \{c \in T : m(c) = L_i\} \\
&\tau_i^{GT} = \max_{c \in L_i} \tau_c \\
&\tau_{min} = \min_c \tau_c \\
&\hat{\tau}_i^{GT} = \begin{cases} \text{median}_{c \in \mathcal{C}_i} \; \tau_c & |\mathcal{C}_i| > 0, \\ \tau_{min} & |\mathcal{C}_i| = 0 \end{cases} \\
&TTS_i = 1 - \frac{\tau_i^{GT} - \hat{\tau}_i^{GT}}{\tau_i^{GT} - \tau_{min}}
\end{aligned}
\tag{4}
$$

- Terminal Temporal Precision (TTP): quantifies the proportion of inferred terminal cells that belong to ground-truth terminal clusters, independently of their temporal position (Equation 5). Values lie in $[0, 1]$, and larger values indicate purer terminal-state composition.

$$
TTP = \frac{|\{c \in T : m(c) \in L\}|}{|T|} \tag{5}
$$

- Terminal State Recall (TSR): quantifies whether the full set of expected terminal populations is recovered (Equation 6). Values lie in $[0, 1]$, and a value of 1 indicates that all ground-truth terminal clusters are represented among the inferred terminal states.

$$
\begin{aligned}
&\hat{L} = \{m(c) : c \in T\} \\
&TSR = \frac{|\hat{L} \cap L|}{|L|}
\end{aligned}
\tag{6}
$$

- Terminal Temporal Concentration (TTC): quantifies how tightly inferred terminal cells are localized along pseudotime within each ground-truth terminal cluster. The final metric averages $TTC_i$ across clusters. Because pseudotime ranges in $[0, 1]$, $TTC_i$ also lies in $[0, 1]$, and larger values indicate greater temporal coherence.

$$
\begin{aligned}
&IQR_i = Q_{0.75}(\{\tau_c | c \in T \cap L_i\}) - Q_{0.25}(\{\tau_c | c \in T \cap L_i\}) \\
&TTC_i = 1 - IQR_i
\end{aligned}
\tag{7}
$$

- Terminal State Score: geometric mean of TTS, TTP, TSR, and TTC.

Unsupervised metrics assess whether fate probabilities are coherently organized across cells and reflect gradual commitment. Collectively, they evaluate four complementary aspects of fate inference:

1. Progressive loss of cellular plasticity during differentiation: as cells commit to a lineage, plasticity is expected to decrease along pseudotime. This is captured by correlating the entropy-based measures with pseudotime, using Spearman and Pearson coefficients. A coherent model should yield negative correlation values between pseudotime and Shannon entropy (decrease in uncertainty as cells differentiate) and a positive correlation between pseudotime and KL divergence, recovering the increasing divergence from the population-average fate profile. The strength and sign of these correlations therefore serve as a quantitative check on whether the inferred fate probabilities are consistent with progressive commitment.

2. Progressive commitment toward specific terminal fates: quantified by the Fate Concentration Index (FCI), defined as the Spearman correlation between pseudotime and fate concentration (Equation 8). As differentiation proceeds, fate probability mass is expected to concentrate on fewer terminal states.

$$\lambda_c := \sum_{k \in \mathcal{T}} F_{c,k}^2 \tag{8}$$

3. Association of inferred terminal states with terminal differentiation stages: quantified by the Terminal Pseudotime Enrichment Score (TPES), which measures pseudotime enrichment within inferred terminal states relative to the dataset-wide pseudotime distribution (Equation 9). Positive scores indicate enrichment toward terminal differentiation, whereas values near zero or below indicate weak or absent temporal localization. A rank-based variant replaces pseudotime values with normalized cell ranks to obtain a scale-invariant version that depends only on relative ordering.

$$\begin{aligned} TPES_k &= \underset{c \in \mathcal{T}_k}{\text{median}}(\tau_c) - \underset{c}{\text{median}}(\tau_c) \\ TPES &= \frac{1}{K} \sum_k TPES_k \end{aligned} \tag{9}$$

4. Resolution and separation of terminal fates: quantified by the terminal-state silhouette metric $S$. Cells associated with the same terminal fate are expected to have similar fate-probability profiles, whereas cells committed to different fates should remain separated in fate-probability space. Two formulations are considered. The first is a pseudotime-weighted silhouette that emphasizes late-stage cells to account for gradual fate commitment (Equation 10).

$$\begin{aligned} \mathcal{T}_c &= \underset{k}{\text{argmax}} \ F_{c,k}, \\ a_c &= \frac{1}{|\mathcal{T}_c| - 1} \sum_{j \in \mathcal{T}_c,\, j \neq c} d(c,j), \\ b_i &= \min_{k \neq \mathcal{T}_c} \left( \frac{1}{|\mathcal{T}_c|} \sum_{j \in \mathcal{T}_c} d(c,j) \right) \\ s_c &= \frac{b_c - a_c}{max(a_c, b_c)} \\ S &= \frac{\sum_c \tau_c^\alpha s_c}{\sum_c \tau_c^\alpha} \end{aligned} \tag{10}$$

The second is a soft silhouette that replaces hard terminal-state assignments with probability-weighted similarities, thereby preserving information about fate uncertainty and avoiding artificial lineage boundaries (Equation 11). Silhouette values lie in $[-1, 1]$, and larger values indicate more coherent and better-resolved terminal outcomes.

$$
\begin{aligned}
w_{c,j} &= \sum_{k=1}^{K} F_{c,k} F_{j,k}, \\
a_c &= \frac{\sum_{j \neq c} w_{c,j} d(c,j)}{\sum_{j \neq c} w_{c,j}}, \\
b_c &= \frac{\sum_{c \neq j} (1 - w_{c,j}) d(c,j)}{\sum_{j \neq i} (1 - w_{c,j})}, \\
s_c &= \frac{b_c - a_c}{\max(a_c, b_c)}, \\
S &= \frac{1}{N} \sum_c s_c
\end{aligned} \tag{11}
$$

### 6 Multimodal Synthetic Data Simulation and Integrated Trajectory Inference with ATLAS

We generated simulated multimodal datasets with scMultiSim v1.0.0 [12]. The simulator produces paired scRNA-seq and scATAC-seq data together with ground-truth pseudotime, lineage assignments, and terminal states.

Each dataset consists of 1,000 cells generated with the `sim_true_counts` function. We used the preconfigured 100-gene Gene Regulatory Network (GRN) provided by the simulator together with two differentiation trees corresponding to topologies with three and five branches (Supplementary Fig. 2 a).

We explored multiple combinations of the parameters $r_d$ and $\sigma_{cif}$ (Supplementary Table 1). These parameters control the dependence of the simulation on the differentiation-tree structure and the degree of within-cluster cellular heterogeneity, respectively. Lower values of $r_d$ and higher values of $\sigma_{cif}$ produce increasingly diffuse trajectories (Supplementary Figure 2 b). This design generated 25 simulated datasets with different degrees of trajectory ambiguity.

We derived gene activity scores from simulated scATAC-seq data by aggregating accessibility across gene-associated regulatory regions. We preprocessed the data with Scanpy v1.10.4 and Muon v0.1.7 [5, 2, 13]. The workflow included normalization, log-transformation of scRNA-seq data, PCA, and construction of the WNN graph integrating gene expression and gene activity [3, 14]. To evaluate the effect of neighborhood size in each modality and in the WNN, we tested multiple values for the number of nearest neighbors used during graph construction (Supplementary Table 1). We retained 20 principal components for gene expression and 10 for gene activity.

For supervised evaluation, we constructed ground-truth fate probabilities from the developmental tree and cluster annotations (Supplementary Fig. 2). For each cluster, fate probabilities were defined uniformly over the set of reachable terminal states. Root clusters were assigned a uniform distribution over all terminal states, parent clusters were assigned a uniform distribution over the reachable subset, and leaf clusters were one-hot encoded.

When required by the supervised metrics, we fixed initial and terminal states to predefined cell sets. We defined initial states as the cells with the lowest ground-truth pseudotime values in the dataset and terminal states as the cells with the highest pseudotime values within each leaf cluster.

We first configured ATLAS to use Palantir as the trajectory-inference backbone. We ran Trajectory Inference (TI) in two configurations: automatic inference of terminal states and explicit specification of ground-truth terminal cells. In both cases, we used the earliest cell as the root state. We set the number of Diffusion Components (DC) to 5 and used 250 waypoints to compute pseudotime and fate probabilities.

We also evaluated ATLAS with the CellRank `PseudotimeKernel`. Ground-truth pseudotime values oriented cell-cell transitions, and the WNN connectivity matrix computed during preprocessing served as the neighborhood graph. During transition-matrix construction, we applied the hard-thresholding scheme to prune edges pointing toward decreasing pseudotime.

We considered two configurations: automatic inference of terminal states from the transition dynamics, allowing overlap between initial and terminal macrostates when supported by the data, and explicit specification of initial and terminal states from ground-truth annotations.

In both configurations, the resulting directed Markov chain was used to compute absorption probabilities toward terminal states.

### 7 Trajectory inference with native Palantir and CellRank on scRNA-seq data

To compare ATLAS against the original Palantir and CellRank frameworks, we also performed scRNA-seq-only TI with the native implementations of both methods across all simulated datasets.

We preprocessed scRNA-seq data as in the multimodal setting. We retained 20 principal components and varied the number of nearest neighbors used for graph construction according to Supplementary Table 1, yielding 200 distinct runs. We retrieved ground-truth pseudotime values, lineage assignments, and terminal states as described in Supplementary Section 6.

For Palantir, we constructed a diffusion kernel from the RNA-based KNN graph and computed diffusion maps retaining 5 diffusion components. We then ran Palantir with 250 waypoints, using the earliest cell as the initial state. We considered two configurations: automatic inference of terminal states and explicit specification of ground-truth terminal cells. We aggregated the resulting fate probabilities at the cluster level by summing probabilities across cells assigned to the same annotated population, thereby obtaining a cell-by-terminal-cluster probability matrix.

For CellRank, we constructed a `PseudotimeKernel` using ground-truth pseudotime as temporal ordering and RNA-based neighborhood-graph connectivities to define local transitions. We computed the transition matrix with the hard-thresholding scheme and inferred macrostates and fate probabilities with the Generalized Perron Cluster Cluster Analysis (GPCCA) estimator. The number of macrostates was not fixed a priori. As for Palantir, we considered automatic inference of initial and terminal states, allowing overlap, and explicit assignment of ground-truth cell subsets.

To maintain consistency with ATLAS, we recomputed cell-fate uncertainty independently of the native framework outputs. For both Palantir and CellRank, we computed Shannon entropy from the inferred fate probability distributions and KL divergence from each cell-specific distribution relative to the population-average fate distribution. We then min–max normalized both quantities to the $[0, 1]$ range to support direct comparison across simulations and methods.

### 8 Experiments on the Fresh Embryonic E18 Mouse Brain

We obtained paired scRNA-seq and scATAC-seq profiles, gene annotations, and fragment files for the Fresh Embryonic E18 Mouse Brain dataset from 10x Genomics [15]. We retrieved cell-type annotations from the MultiVelo repository [16].

For scRNA-seq, we applied Median Absolute Deviation (MAD)-based quality control as described in [17] and excluded cells with mitochondrial transcript content greater than 10%. We then log-normalized the expression matrix and selected the top 2,000 highly variable genes. For scATAC-seq, we discarded cells with nucleosome signal greater than 2 and Transcripton Starting Site (TSS) enrichment lower than 1.5.

We restricted the dataset to cortical populations (”Upper Layer”, ”Deeper Layer”, ”Subplate”, ”V-SVZ”), progenitor populations (”RG, Astro, OPC”), and the structural cell type ”Ependymal”.

We filtered gene coordinates to retain canonical chromosomes (1–19, X, and Y) and provided them to ATLAS for gene activity computation from scATAC-seq data. We performed PCA on both expression and activity modalities with Scanpy default settings.

ATLAS then constructed the single-modality KNN graphs and the WNN graph. Based on PCA variance-plot inspection, we retained 20 PCs for gene expression and 10 for gene activity. We computed single-modality KNN graphs with 15 neighbors and set the number of neighbors for WNN construction to `None`, allowing the method to use the arithmetic mean of modality-specific neighborhood sizes.

We performed trajectory inference with ATLAS using both Palantir- and CellRank-based strategies. In the Palantir-based configuration, we selected a random cell from the ”RG, Astro, OPC” population as the initial state and used default parameters. We then used the inferred pseudotime within CellRank to orient the transition matrix. We ran the analysis with six and twelve macrostates. The lower setting reflects prior biological expectations, whereas the higher-resolution setting tests how increasing the number of available macrostates affects branch construction and the partitioning of initial, intermediate, and terminal states.

For comparison, we applied standalone CellRank and Palantir to the gene expression-derived KNN graph using the same parameter settings and initial cell.

### 9 Experiments on the SHARE-seq Mouse Hair Follicle

We obtained multi-omics data for late anagen mouse skin from the original study [18]. We selected cells involved in Hair Shaft (HS) formation based on the original annotations, retaining Transient Amplifying Cells (TAC) (TAC-1 and TAC-2), Medulla, Inner Root Cells (IRS), and HS Cortex populations.

For scRNA-seq, we applied the same MAD-based filtering strategy used In Supplementary Section 8 [17] and excluded cells with mitochondrial transcript content above 10%. We log-normalized the filtered counts and selected the top 2,000 highly variable genes. For scATAC-seq, we excluded cells with nucleosome signal greater than 2.

We retrieved gene coordinates from the mm10 reference genome with `pybiomart` [19] and restricted them to canonical chromosomes (1–19 and X) for gene activity computation.

We computed gene activity scores with the `compute_fragments_features` function from muon, setting the `count_reads` parameter to `False` to match the BED4 fragment-file format, in which each fragment contributes a single count. We then normalized the gene activity values.

We performed PCA on each modality independently with Scanpy default parameters.

ATLAS then constructed modality-specific KNN graphs and the WNN graph, retaining 20 PCs for gene expression and 10 for gene activity, with $K = 15$ neighbors. We performed TI with both available ATLAS strategies, using a randomly selected ”TAC-1” cell as the initial state when required. We ran the analysis with four and eight macrostates to assess how increasing the number of states affects branch construction.

For comparison, we also ran standalone CellRank and Palantir on the gene expression-derived KNN graph with the same parameter settings and initial cell.

### 10 Experiments on the Human Fetal Brain

The Human Fetal Brain dataset [20] contains paired gene-expression and gene-activity values for approximately 8,000 cells and was already filtered according to the quality criteria reported in the original publication. Cell-cluster labels are provided with the count matrices, and we renamed the original clusters according to the Multivelo guidelines [16].

We log-normalized the scRNA-seq matrix and selected the 2,000 most differentially expressed genes. We normalized gene-activity values. We computed PCA for both modalities with Scanpy default parameters and retained 20 components for gene expression and 10 for gene activity during unimodal KNN graph construction ($K = 15$). We then built the WNN graph with ATLAS using default settings.

We performed TI with both available ATLAS strategies, using a randomly selected Cycling Progenitor Cell as the initial state when required. We ran the analysis with five and ten macrostates to assess how increasing the number of available states affects branch construction and the delineation of initial and terminal populations.

For comparison, we also ran standalone CellRank and Palantir on the gene expression-derived KNN graph with the same parameter settings and initial cell.

### 11 Evaluation of ATLAS against scRNA-seq only experiments

We compared ATLAS against the original Palantir and CellRank methods on the synthetic datasets using scRNA-seq-only experiments as baselines. Results are reported in Supplementary Tables 3 and 4.

We performed comparisons separately for each dataset condition defined by tree topology and terminal-constraint setting. For each method, we executed multiple runs across the shared hyperparameter grid in Supplementary Table 1. We evaluated performance at the level of hyperparameter configurations to ensure comparability under identical experimental conditions.

Some runs yielded undefined metric values because the inferred structures were degenerate or the metric could not be computed under a given configuration. To avoid optimistic bias from excluding such cases, we assigned undefined values the worst attainable score $w$ for the corresponding metric. Formally, the penalized metric is defined as:

$$\tilde{m} = \begin{cases} m & \text{if defined} \\ w & \text{if undefined} \end{cases} \tag{12}$$

This penalization incorporates degenerate solutions into the performance estimate. Consequently, the summaries reflect expected utility over the full configuration space rather than only over successful runs.

We then grouped runs by tree topology, terminal-constraint setting, and shared hyperparameter configuration. Within each group, we computed the mean penalized metric for each algorithm and retained only configurations present for both methods.

The number of runs per configuration was not identical across methods because ATLAS includes two additional hyperparameters. A direct comparison of global averages would therefore weight configurations by run availability and confound sampling density with algorithm performance. To avoid this bias, we adopted a paired comparison strategy. We retained only hyperparameter configurations shared by ATLAS and the scRNA-seq-only experiments. For each shared configuration $i$, we defined the performance difference as:

$$\Delta_i = \tilde{m}_{ATLAS,i} - \tilde{m}_{Method,i} \tag{13}$$

where $\tilde{m}_{ATLAS,i}$ and $\tilde{m}_{Method,i}$ denote the mean penalized metric across runs for that configuration.

For each dataset, we summarized the distribution of $\Delta_i$ across configurations with both the mean and the median to measure average performance differences between ATLAS and the original scRNA-seq-based frameworks. We also estimated the empirical probability $\hat{p} = P(\Delta_i > 0)$ to quantify the fraction of configurations in which ATLAS outperforms the baseline.

We quantified uncertainty with non-parametric bootstrap resampling over hyperparameter configurations and computed percentile-based 95% confidence intervals. For the probability estimate, we used the Wilson score interval for binomial proportions.

The primary reported quantities represent the expected difference in penalized performance across the shared configuration space.

### 12 Supplementary Tables

| Parameter | Values |
|---|---|
| $\sigma_{cif}$ | [0.1, 0.3, 0.5, 0.7, 0.9] |
| $r_d$ | [0.1, 0.3, 0.5, 0.7, 0.9] |
| KNN (gene expression) | [10, 30, 50, 70] |
| KNN (gene activity) | [10, 30, 50, 70] |
| KNN (WNN) | [10, 30, 50, 70] |

**Supplementary Table 1.** Parameter values explored during synthetic data generation and preprocessing.

| | ATLAS vs Palantir | | ATLAS vs CellRank | |
|---|---|---|---|---|
| | ATLAS | Baseline | ATLAS | Baseline |
| **Five Branches** | | | | |
| Total number of runs | 1,600 | 100 | 1,600 | 100 |
| Total number of valid runs | 1,600 | 100 | 1,304 | 68 |
| Proportion of valid runs | 1.000 (0.963–1.000) | 1.000 (0.963–1.000) | 0.815 (0.759–0.833) | 0.680 (0.583–0.763) |
| **Three Branches** | | | | |
| Total number of runs | 1,600 | 100 | 1,600 | 100 |
| Total number of valid runs | 1,600 | 100 | 1,334 | 88 |
| Proportion of valid runs | 1.000 (0.963–1.000) | 1.000 (0.963–1.000) | 0.833 (0.815–0.851) | 0.880 (0.802–0.930) |

**Supplementary Table 2.** Proportion of correctly ended simulations on synthetic data. Confidence intervals are reported in parentheses.

**(a)** Three-branches synthetic dataset.

| | Terminal States Automatically Inferred | | | Terminal States Fixed | | |
|---|---|---|---|---|---|---|
| Metric | Delta mean (95% CI) | Delta median (95% CI) | P(ATLAS > baseline) | Delta mean (95% CI) | Delta median (95% CI) | P(ATLAS > baseline) |
| Spearman correlation (pseudotime) | 0.018 (-0.034, 0.016) | 0.027 (-0.001, 0.026) | 0.590 (0.492, 0.681) | - | - | - |
| Kendall correlation (pseudotime) | 0.051 (0.007, 0.051) | 0.037 (0.018, 0.037) | 0.640 (0.542, 0.727) | - | - | - |
| Terminal State Score | 0.085 (0.024, 0.082) | 0.086 (0.026, 0.084) | 0.640 (0.542, 0.727) | - | - | - |
| Pearson correlation (Shannon entropy) | -0.343 (-0.535, -0.348) | -0.130 (-0.420, -0.139) | 0.310 (0.228, 0.406) | - | - | - |
| Spearman correlation (Shannon entropy) | -0.343 (-0.530, -0.349) | -0.136 (-0.414, -0.150) | 0.320 (0.237, 0.417) | - | - | - |
| Pearson correlation (KL-divergence) | 0.367 (0.175, 0.362) | 0.165 (0.097, 0.165) | 0.670 (0.573, 0.754) | - | - | - |
| Spearman correlation (KL-divergence) | 0.381 (0.206, 0.378) | 0.190 (0.111, 0.188) | 0.680 (0.583, 0.763) | - | - | - |
| Fate concentration index | 0.155 (0.020, 0.153) | 0.064 (-0.077, 0.056) | 0.540 (0.443, 0.634) | - | - | - |
| Terminal Pseudotime Enrichment Score | 0.033 (-0.063, 0.027) | -0.046 (-0.058, -0.047) | 0.280 (0.201, 0.375) | - | - | - |
| Terminal State Silhouette (soft) | 0.121 (0.024, 0.117) | 0.077 (-0.021, 0.076) | 0.560 (0.462, 0.653) | - | - | - |
| Terminal State Silhouette (pseudotime) | 0.190 (0.063, 0.186) | 0.073 (-0.045, 0.073) | 0.520 (0.423, 0.615) | - | - | - |
| JSD – leaf | - | - | - | -0.027 (-0.038, -0.027) | -0.037 (-0.045, -0.037) | 0.270 (0.193, 0.364) |
| JSD – parent | - | - | - | - | - | - |
| JSD – root | - | - | - | -0.050 (-0.070, -0.050) | -0.047 (-0.060, -0.047) | 0.330 (0.246, 0.427) |

**(b)** Five-branches synthetic dataset.

| | Terminal States Automatically Inferred | | | Terminal States Fixed | | |
|---|---|---|---|---|---|---|
| Metric | Delta mean (95% CI) | Delta median (95% CI) | P(ATLAS > baseline) | Delta mean (95% CI) | Delta median (95% CI) | P(ATLAS > baseline) |
| Spearman correlation (pseudotime) | 0.152 (0.087, 0.148) | 0.085 (0.023, 0.083) | 0.670 (0.573, 0.754) | - | - | - |
| Kendall correlation (pseudotime) | 0.126 (0.071, 0.122) | 0.094 (0.036, 0.091) | 0.670 (0.573, 0.754) | - | - | - |
| Terminal State Score | 0.105 (0.057, 0.103) | 0.034 (-0.009, 0.030) | 0.550 (0.452, 0.644) | - | - | - |
| Pearson correlation (Shannon entropy) | -0.396 (-0.562, -0.403) | -0.107 (-0.486, -0.112) | 0.380 (0.291, 0.478) | - | - | - |
| Spearman correlation (Shannon entropy) | -0.391 (-0.558, -0.397) | -0.082 (-0.451, -0.083) | 0.390 (0.300, 0.488) | - | - | - |
| Pearson correlation (KL-divergence) | 0.326 (0.160, 0.320) | 0.099 (0.000, 0.097) | 0.580 (0.482, 0.672) | - | - | - |
| Spearman correlation (KL-divergence) | 0.332 (0.173, 0.329) | 0.099 (0.000, 0.099) | 0.570 (0.472, 0.663) | - | - | - |
| Fate concentration index | 0.253 (0.109, 0.247) | 0.018 (-0.058, 0.018) | 0.530 (0.433, 0.625) | - | - | - |
| Terminal State Silhouette (soft) | 0.189 (0.092, 0.186) | 0.056 (-0.021, 0.056) | 0.560 (0.462, 0.653) | - | - | - |
| Terminal State Silhouette (pseudotime) | 0.239 (0.125, 0.236) | 0.021 (-0.026, 0.021) | 0.510 (0.413, 0.606) | - | - | - |
| Terminal Pseudotime Enrichment Score | 0.189 (0.095, 0.183) | 0.009 (-0.006, 0.007) | 0.560 (0.462, 0.653) | - | - | - |
| JSD – leaf | - | - | - | -0.002 (-0.018, -0.002) | 0.006 (-0.014, 0.006) | 0.530 (0.433, 0.625) |
| JSD – parent | - | - | - | -0.043 (-0.058, -0.044) | -0.029 (-0.047, -0.029) | 0.260 (0.184, 0.354) |
| JSD – root | - | - | - | -0.064 (-0.088, -0.065) | -0.071 (-0.087, -0.071) | 0.260 (0.184, 0.354) |

**Supplementary Table 3.** ATLAS benchmarking against Palantir on scRNA-seq only data.

**(a)** Three-branches synthetic dataset.

| | Terminal States Automatically Inferred | | | Terminal States Fixed | | |
|---|---|---|---|---|---|---|
| Metric | Delta mean (95% CI) | Delta median (95% CI) | P(ATLAS > baseline) | Delta mean (95% CI) | Delta median (95% CI) | P(ATLAS > baseline) |
| Spearman correlation (pseudotime) | - | - | - | - | - | - |
| Kendall correlation (pseudotime) | - | - | - | - | - | - |
| Terminal State Score | -0.037 (-0.094, -0.038) | -0.010 (-0.016, -0.011) | 0.340 (0.255, 0.437) | - | - | - |
| Pearson correlation (Shannon entropy) | 0.068 (-0.033, 0.066) | 0.024 (0.002, 0.019) | 0.620 (0.522, 0.709) | - | - | - |
| Spearman correlation (Shannon entropy) | 0.078 (-0.033, 0.078) | 0.057 (-0.001, 0.057) | 0.600 (0.502, 0.691) | - | - | - |
| Pearson correlation (KL-divergence) | 0.021 (0.007, 0.021) | 0.021 (0.012, 0.020) | 0.750 (0.657, 0.825) | - | - | - |
| Spearman correlation (KL-divergence) | -0.056 (-0.165, -0.061) | -0.022 (-0.084, -0.022) | 0.410 (0.319, 0.508) | - | - | - |
| Fate concentration index | -0.053 (-0.147, -0.057) | -0.037 (-0.079, -0.037) | 0.440 (0.347, 0.538) | - | - | - |
| Terminal Pseudotime Enrichment Score | -0.063 (-0.147, -0.064) | -0.024 (-0.044, -0.024) | 0.360 (0.273, 0.458) | - | - | - |
| Terminal State Silhouette (soft) | -0.129 (-0.225, -0.132) | -0.098 (-0.198, -0.100) | 0.310 (0.228, 0.406) | - | - | - |
| Terminal State Silhouette (pseudotime) | -0.113 (-0.217, -0.115) | -0.065 (-0.141, -0.065) | 0.320 (0.237, 0.417) | - | - | - |
| JSD – leaf | - | - | - | 0.020 (0.011, 0.020) | 0.010 (0.006, 0.010) | 0.640 (0.542, 0.727) |
| JSD – parent | - | - | - | - | - | - |
| JSD – root | - | - | - | -0.034 (-0.044, -0.034) | -0.036 (-0.047, -0.037) | 0.350 (0.264, 0.447) |

**(b)** Five-branches synthetic dataset.

| | Terminal States Automatically Inferred | | | Terminal States Fixed | | |
|---|---|---|---|---|---|---|
| Metric | Delta mean (95% CI) | Delta median (95% CI) | P(ATLAS > baseline) | Delta mean (95% CI) | Delta median (95% CI) | P(ATLAS > baseline) |
| Spearman correlation (pseudotime) | - | - | - | - | - | - |
| Kendall correlation (pseudotime) | - | - | - | - | - | - |
| Terminal State Score | 0.145 (0.080, 0.144) | 0.027 (-0.013, 0.027) | 0.560 (0.462, 0.653) | - | - | - |
| Pearson correlation (Shannon entropy) | -0.238 (-0.348, -0.242) | -0.040 (-0.277, -0.040) | 0.420 (0.328, 0.518) | - | - | - |
| Spearman correlation (Shannon entropy) | -0.239 (-0.353, -0.242) | -0.012 (-0.268, -0.012) | 0.470 (0.375, 0.567) | - | - | - |
| Pearson correlation (KL-divergence) | 0.309 (0.195, 0.306) | 0.084 (0.016, 0.083) | 0.630 (0.532, 0.718) | - | - | - |
| Spearman correlation (KL-divergence) | 0.284 (0.170, 0.278) | 0.044 (-0.018, 0.044) | 0.570 (0.472, 0.663) | - | - | - |
| Fate concentration index | 0.240 (0.134, 0.232) | 0.004 (-0.030, 0.002) | 0.500 (0.404, 0.596) | - | - | - |
| Terminal Pseudotime Enrichment Score | 0.194 (0.103, 0.190) | 0.015 (-0.015, 0.015) | 0.560 (0.462, 0.653) | - | - | - |
| Terminal State Silhouette (soft) | 0.197 (0.084, 0.194) | -0.028 (-0.087, -0.032) | 0.460 (0.366, 0.557) | - | - | - |
| Terminal State Silhouette (pseudotime) | 0.219 (0.097, 0.213) | 0.219 (0.097, 0.213) | 0.480 (0.385, 0.577) | - | - | - |
| JSD – leaf | - | - | - | 0.018 (0.010, 0.018) | 0.015 (0.002, 0.015) | 0.600 (0.502, 0.691) |
| JSD – parent | - | - | - | -0.035 (-0.047, -0.036) | -0.035 (-0.047, -0.035) | 0.260 (0.184, 0.354) |
| JSD – root | - | - | - | -0.073 (-0.084, -0.074) | -0.064 (-0.077, -0.064) | 0.070 (0.034, 0.137) |

**Supplementary Table 4.** ATLAS benchmarking against CellRank on scRNA-seq only data.

**(a)** SHARE-seq mouse hair follicle ATLAS and Palantir applied on scRNA-seq data.

| | ATLAS | | scRNA-seq only | |
|---|---|---|---|---|
| Metric Name | Statistic | CI [LL, UL] | Statistic | CI [LL, UL] |
| Pearson correlation (pseudotime – Shannon entropy) | -0.912 | [-0.916, -0.908] | -0.919 | [-0.923, -0.916] |
| Pearson correlation (pseudotime – KL divergence) | 0.826 | [0.818, 0.834] | 0.792 | [0.783, 0.801] |
| Spearman correlation (pseudotime – Shannon entropy) | -0.838 | [-0.848, -0.828] | -0.759 | [-0.773, -0.744] |
| Spearman correlation (pseudotime – KL divergence) | 0.896 | [0.890, 0.901] | 0.888 | [0.883, 0.893] |
| Fate Concentration Index | 0.838 | [0.828, 0.848] | 0.759 | [0.744, 0.773] |
| Terminal Pseudotime Enrichment Score | 0.483 | - | 0.467 | - |
| Terminal State Silhouette (pseudotime) | 0.576 | - | 0.726 | - |
| Terminal State Silhouette (soft) | 0.215 | - | 0.266 | - |

**(b)** SHARE-seq mouse hair follicle ATLAS and CellRank `PseudotimeKernel` applied on scRNA-seq data (4 macrostates).

| | ATLAS | | scRNA-seq only | |
|---|---|---|---|---|
| Metric Name | Statistic | CI [LL, UL] | Statistic | CI [LL, UL] |
| Pearson correlation (pseudotime – Shannon entropy) | -0.848 | [-0.855, -0.841] | -0.922 | [-0.925, -0.918] |
| Pearson correlation (pseudotime – KL divergence) | 0.827 | [0.820, 0.835] | 0.899 | [0.895, 0.904] |
| Spearman correlation (pseudotime – Shannon entropy) | -0.847 | [-0.855, -0.839] | -0.876 | [-0.884, -0.868] |
| Spearman correlation (pseudotime – KL divergence) | 0.902 | [0.896, 0.907] | 0.901 | [0.895, 0.906] |
| Fate Concentration Index | 0.851 | [0.843, 0.859] | 0.875 | [0.867, 0.882] |
| Terminal Pseudotime Enrichment Score | 0.445 | - | 0.457 | - |
| Terminal State Silhouette (pseudotime) | 0.371 | - | 0.464 | - |
| Terminal State Silhouette (soft) | 0.121 | - | 0.136 | - |

**(c)** SHARE-seq mouse hair follicle ATLAS and CellRank `PseudotimeKernel` applied on scRNA-seq data (8 macrostates).

| | ATLAS | | scRNA-seq only | |
|---|---|---|---|---|
| Metric Name | Statistic | CI [LL, UL] | Statistic | CI [LL, UL] |
| Pearson correlation (pseudotime – Shannon entropy) | -0.825 | [-0.832, -0.817] | -0.874 | [-0.880, -0.868] |
| Pearson correlation (pseudotime – KL divergence) | 0.794 | [0.785, 0.803] | 0.664 | [0.650, 0.677] |
| Spearman correlation (pseudotime – Shannon entropy) | -0.853 | [-0.861, -0.845] | -0.878 | [-0.886, -0.869] |
| Spearman correlation (pseudotime – KL divergence) | 0.892 | [0.887, 0.898] | 0.831 | [0.820, 0.842] |
| Fate Concentration Index | 0.854 | [0.846, 0.862] | 0.854 | [0.845, 0.863] |
| Terminal Pseudotime Enrichment Score | 0.439 | - | 0.266 | - |
| Terminal State Silhouette (pseudotime) | 0.411 | - | 0.426 | - |
| Terminal State Silhouette (soft) | 0.116 | - | 0.136 | - |

**Supplementary Table 5.** Unsupervised benchmarking results comparing ATLAS and the original frameworks applied on scRNA-seq data only.

**(a)** Fresh Embryonic E18 Mouse Brain Dataset ATLAS and Palantir applied on scRNA-seq data.

| | ATLAS | | scRNA-seq only | |
|---|---|---|---|---|
| Metric Name | Statistic | CI [LL, UL] | Statistic | CI [LL, UL] |
| Pearson correlation (pseudotime – Shannon entropy) | - | - | - | - |
| Pearson correlation (pseudotime – KL divergence) | - | - | - | - |
| Spearman correlation (pseudotime – Shannon entropy) | - | - | - | - |
| Spearman correlation (pseudotime – KL divergence) | - | - | - | - |
| Fate Concentration Index | -0.339 | [-0.372, -0.305] | 0.708 | [0.686, 0.730] |
| Terminal Pseudotime Enrichment Score | 0.244 | - | 0.5 | - |
| Terminal State Silhouette (pseudotime) | - | - | - | - |
| Terminal State Silhouette (soft) | - | - | - | - |

**(b)** Fresh Embryonic E18 Mouse Brain Dataset ATLAS and CellRank `PseudotimeKernel` applied on scRNA-seq data (6 macrostates).

| | ATLAS | | scRNA-seq only | |
|---|---|---|---|---|
| Metric Name | Statistic | CI [LL, UL] | Statistic | CI [LL, UL] |
| Pearson correlation (pseudotime – Shannon entropy) | -0.643 | [-0.662, -0.624] | -0.792 | [-0.804, -0.780] |
| Pearson correlation (pseudotime – KL divergence) | 0.676 | [0.657, 0.693] | -0.286 | [-0.316, -0.256] |
| Spearman correlation (pseudotime – Shannon entropy) | -0.608 | [-0.633, -0.583] | -0.683 | [-0.710, -0.655] |
| Spearman correlation (pseudotime – KL divergence) | 0.648 | [0.624, 0.671] | 0.174 | [0.132, 0.215] |
| Fate Concentration Index | 0.599 | [0.573, 0.624] | 0.671 | [0.643, 0.698] |
| Terminal Pseudotime Enrichment Score | 0.371 | - | -0.104 | - |
| Terminal State Silhouette (pseudotime) | 0.513 | - | 0.821 | - |
| Terminal State Silhouette (soft) | 0.245 | - | 0.618 | - |

**(c)** Fresh Embryonic E18 Mouse Brain Dataset ATLAS and CellRank `PseudotimeKernel` applied on scRNA-seq data (12 macrostates).

| | ATLAS | | scRNA-seq only | |
|---|---|---|---|---|
| Metric Name | Statistic | CI [LL, UL] | Statistic | CI [LL, UL] |
| Pearson correlation (pseudotime – Shannon entropy) | -0.596 | [-0.617, -0.574] | -0.649 | [-0.667, -0.629] |
| Pearson correlation (pseudotime – KL divergence) | 0.663 | [0.644, 0.681] | -0.080 | [-0.113, -0.047] |
| Spearman correlation (pseudotime – Shannon entropy) | -0.514 | [-0.543, -0.483] | -0.713 | [-0.739, -0.686] |
| Spearman correlation (pseudotime – KL divergence) | 0.561 | [0.532, 0.589] | 0.081 | [0.046, 0.116] |
| Fate Concentration Index | 0.509 | [0.478, 0.538] | 0.702 | [0.675, 0.729] |
| Terminal Pseudotime Enrichment Score | 0.458 | - | -0.036 | - |
| Terminal State Silhouette (pseudotime) | 0.581 | - | 0.528 | - |
| Terminal State Silhouette (soft) | 0.271 | - | 0.414 | - |

**Supplementary Table 6.** Unsupervised benchmarking results comparing ATLAS and the original frameworks applied on scRNA-seq data only.

**(a)** Fetal Human Brain Dataset ATLAS and Palantir applied on scRNA-seq data.

| | ATLAS | | scRNA-seq only | |
|---|---|---|---|---|
| Metric Name | Statistic | CI [LL, UL] | Statistic | CI [LL, UL] |
| Pearson correlation (pseudotime – Shannon entropy) | - | - | - | - |
| Pearson correlation (pseudotime – KL divergence) | 0.595 | [0.580, 0.610] | - | - |
| Spearman correlation (pseudotime – Shannon entropy) | - | - | - | - |
| Spearman correlation (pseudotime – KL divergence) | 0.144 | [0.123, 0.163] | - | - |
| Fate Concentration Index | 0.080 | [0.052, 0.109] | 0.145 | [0.125, 0.166] |
| Terminal Pseudotime Enrichment Score | 0.401 | - | 0.500 | - |
| Terminal State Silhouette (pseudotime) | 0.999 | - | - | - |
| Terminal State Silhouette (soft) | 0.999 | - | - | - |

**(b)** Fetal Human Brain Dataset ATLAS and CellRank `PseudotimeKernel` applied on scRNA-seq data (5 macrostates).

| | ATLAS | | scRNA-seq only | |
|---|---|---|---|---|
| Metric Name | Statistic | CI [LL, UL] | Statistic | CI [LL, UL] |
| Pearson correlation (pseudotime – Shannon entropy) | -0.819 | [-0.827, -0.811] | -0.470 | [-0.488, -0.451] |
| Pearson correlation (pseudotime – KL divergence) | 0.720 | [0.709, 0.732] | 0.575 | [0.559, 0.591] |
| Spearman correlation (pseudotime – Shannon entropy) | -0.916 | [-0.922, -0.909] | -0.503 | [-0.524, -0.481] |
| Spearman correlation (pseudotime – KL divergence) | 0.792 | [0.780, 0.804] | 0.387 | [0.362, 0.411] |
| Fate Concentration Index | 0.895 | [0.889, 0.902] | 0.459 | [0.436, 0.481] |
| Terminal Pseudotime Enrichment Score | 0.397 | - | 0.373 | - |
| Terminal State Silhouette (pseudotime) | 0.635 | - | 0.511 | - |
| Terminal State Silhouette (soft) | 0.431 | - | 0.368 | - |

**(c)** Fetal Human Brain Dataset Dataset ATLAS and CellRank `PseudotimeKernel` applied on scRNA-seq data (10 macrostates).

| | ATLAS | | scRNA-seq only | |
|---|---|---|---|---|
| Metric Name | Statistic | CI [LL, UL] | Statistic | CI [LL, UL] |
| Pearson correlation (pseudotime – Shannon entropy) | -0.802 | [-0.810, -0.793] | -0.470 | [-0.488, -0.451] |
| Pearson correlation (pseudotime – KL divergence) | 0.782 | [0.772, 0.791] | 0.570 | [0.554, 0.586] |
| Spearman correlation (pseudotime – Shannon entropy) | -0.907 | [-0.915, -0.898] | -0.583 | [-0.601, -0.565] |
| Spearman correlation (pseudotime – KL divergence) | 0.764 | [0.750, 0.777] | 0.501 | [0.477, 0.524] |
| Fate Concentration Index | 0.884 | [0.874, 0.893] | 0.531 | [0.512, 0.551] |
| Terminal Pseudotime Enrichment Score | 0.324 | - | 0.258 | - |
| Terminal State Silhouette (pseudotime) | 0.602 | - | 0.791 | - |
| Terminal State Silhouette (soft) | 0.353 | - | 0.638 | - |

**Supplementary Table 7.** Unsupervised benchmarking results comparing ATLAS and the original frameworks applied on scRNA-seq data only.

**(a)** Fetal Human Brain Dataset ATLAS and Palantir applied on scRNA-seq data.

| | ATLAS | | scRNA-seq only | |
|---|---|---|---|---|
| Metric Name | Statistic | CI [LL, UL] | Statistic | CI [LL, UL] |
| Pearson correlation (pseudotime – Shannon entropy) | -0.605 | [-0.620, -0.590] | -0.110 | [0.133, -0.087] |
| Pearson correlation (pseudotime – KL divergence) | 0.709 | [0.697, 0.720] | 0.392 | [0.371, 0.411] |
| Spearman correlation (pseudotime – Shannon entropy) | -0.551 | [-0.578, -0.523] | 0.127 | [0.095, 0.157] |
| Spearman correlation (pseudotime – KL divergence) | 0.714 | [0.692, 0.735] | 0.163 | [0.133, 0.193] |
| Fate Concentration Index | 0.597 | [0.570, 0.621] | -0.176 | [-0.206, 0.145] |
| Terminal Pseudotime Enrichment Score | 0.426 | - | 0.165 | - |
| Terminal State Silhouette (pseudotime) | 0.557 | - | 0.544 | - |
| Terminal State Silhouette (soft) | 0.189 | - | 0.114 | - |

**(b)** Fetal Human Brain Dataset ATLAS and CellRank `PseudotimeKernel` applied on scRNA-seq data (5 macrostates).

| | ATLAS | | scRNA-seq only | |
|---|---|---|---|---|
| Metric Name | Statistic | CI [LL, UL] | Statistic | CI [LL, UL] |
| Pearson correlation (pseudotime – Shannon entropy) | -0.785 | [-0.794, -0.776] | -0.311 | [-0.322, -0.290] |
| Pearson correlation (pseudotime – KL divergence) | 0.851 | [0.845, 0.858] | 0.510 | [0.492, 0.527] |
| Spearman correlation (pseudotime – Shannon entropy) | -0.831 | [-0.840, -0.821] | -0.541 | [-0.563, -0.517] |
| Spearman correlation (pseudotime – KL divergence) | 0.894 | [0.887, 0.900] | 0.506 | [0.485, 0.528] |
| Fate Concentration Index | 0.795 | [0.783, 0.806] | 0.516 | [0.492, 0.539] |
| Terminal Pseudotime Enrichment Score | 0.423 | - | 0.397 | - |
| Terminal State Silhouette (pseudotime) | 0.507 | - | 0.764 | - |
| Terminal State Silhouette (soft) | 0.301 | - | 0.606 | - |

**(c)** Fetal Human Brain Dataset Dataset ATLAS and CellRank `PseudotimeKernel` applied on scRNA-seq data (10 macrostates).

| | ATLAS | | scRNA-seq only | |
|---|---|---|---|---|
| Metric Name | Statistic | CI [LL, UL] | Statistic | CI [LL, UL] |
| Pearson correlation (pseudotime – Shannon entropy) | -0.820 | [-0.828, -0.812] | -0.577 | [-0.592, -0.561] |
| Pearson correlation (pseudotime – KL divergence) | 0.931 | [0.927, 0.934] | 0.506 | [0.488, 0.523] |
| Spearman correlation (pseudotime – Shannon entropy) | -0.823 | [-0.830, -0.815] | -0.624 | [-0.642, -0.607] |
| Spearman correlation (pseudotime – KL divergence) | 0.939 | [0.935, 0.943] | 0.451 | [0.427, 0.473] |
| Fate Concentration Index | 0.799 | [0.790, 0.808] | 0.590 | [0.571, 0.608] |
| Terminal Pseudotime Enrichment Score | 0.380 | - | 0.209 | - |
| Terminal State Silhouette (pseudotime) | 0.562 | - | 0.735 | - |
| Terminal State Silhouette (soft) | 0.294 | - | 0.597 | - |

**Supplementary Table 8.** Unsupervised benchmarking results comparing ATLAS and the original frameworks applied on scRNA-seq data only (REMOVED OUTLIERS).

### 13 Supplementary Figures

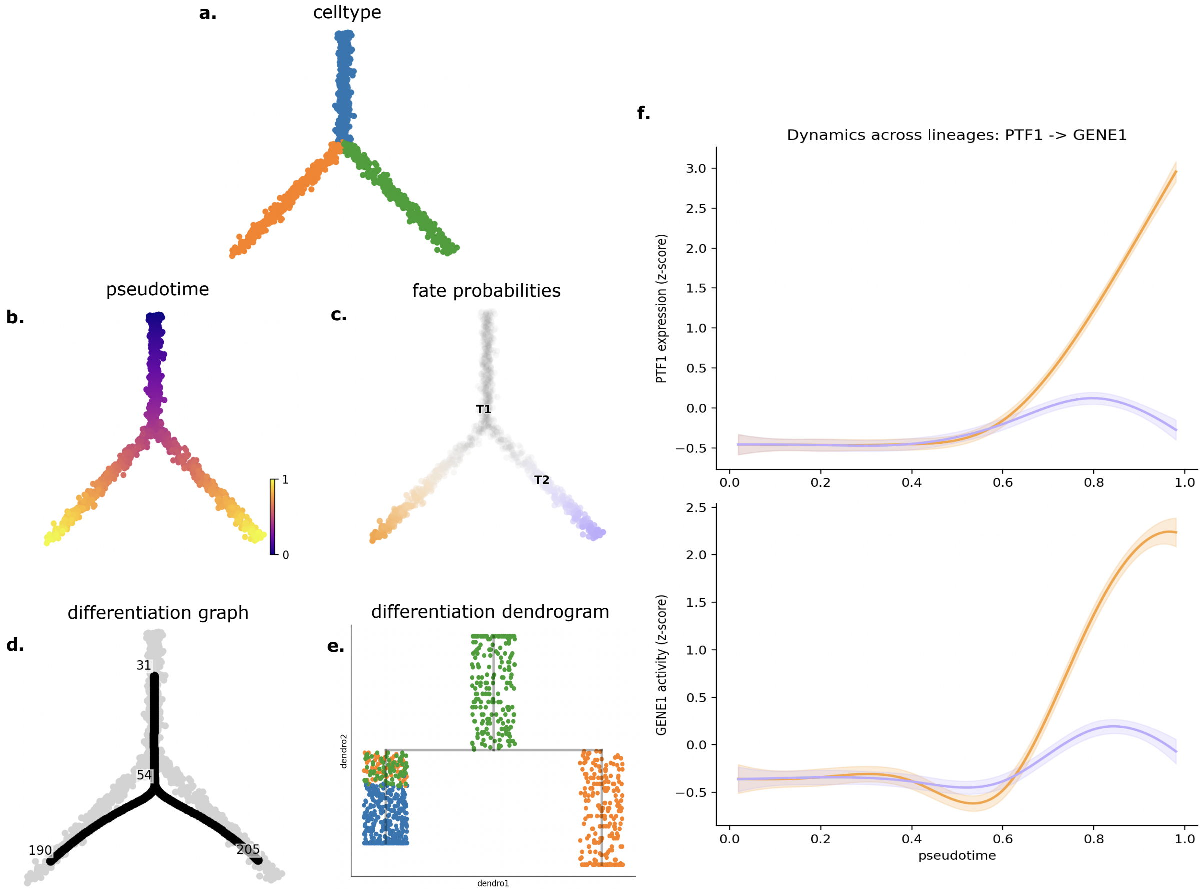


**Supplementary Figure 1.** Example of visualizations provided by ATLAS on a mock dataset. **(a)** UMAP plot, cells colored according to cluster. **(b)** UMAP plot, cells colored according to pseudotime. Similar visualization can be performed with entropy values. **(c)** UMAP plot, cells colored according to most probable fate. **(d)** Principal Graph on UMAP. **(e)** Dendrogram representing the learned principal graph, cells colored according to cluster. **(f)** Gene expression for a transcription factor (on the top) and gene activity for a putative target gene (on the bottom).

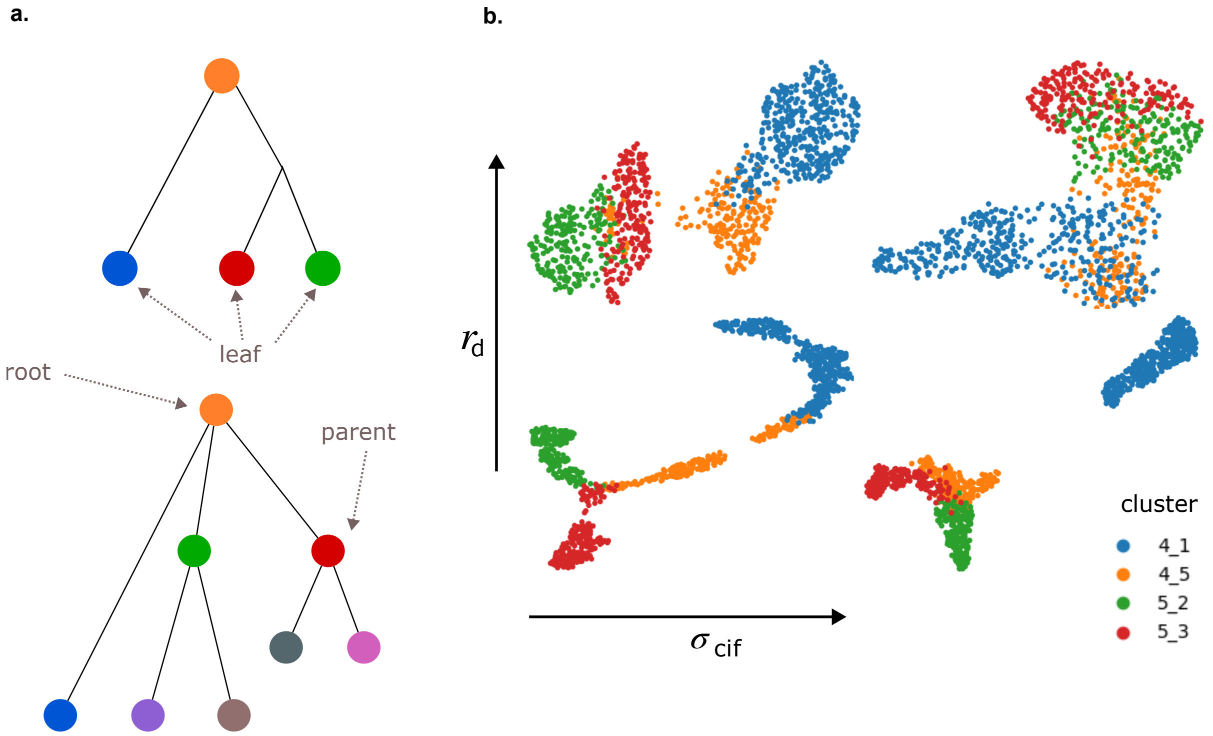


**Supplementary Figure 2.** **a.** Schematic representation of the developmental trees provided as input to scMultiSim and annotation of clusters as root, parent and leaf. **b.** UMAP visualization of the same dataset for increasing values of $r_d$ and $\sigma_{cif}$.

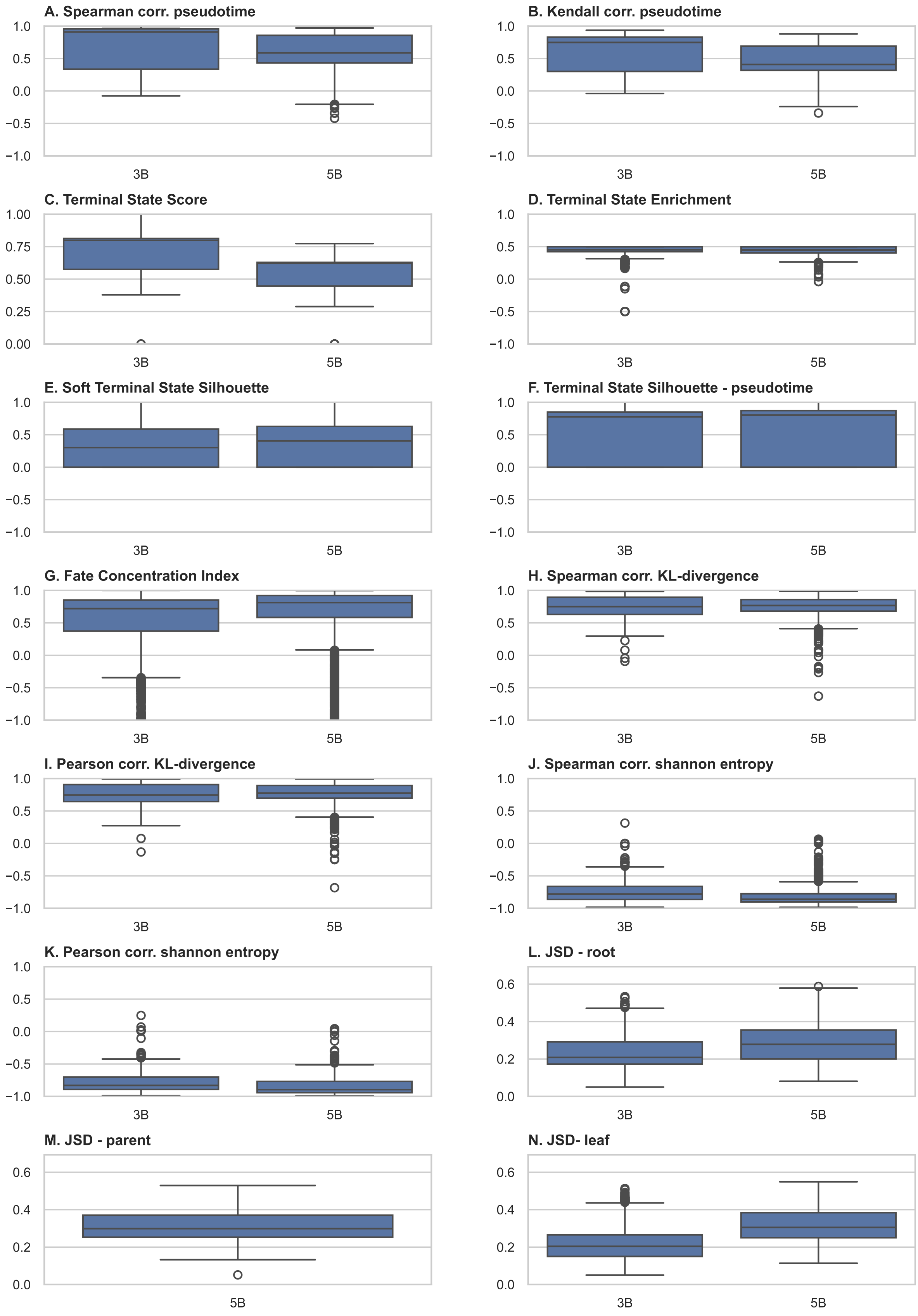


**Supplementary Figure 3.** Evaluation of ATLAS performance with Palantir as TI strategy on the synthetic data.

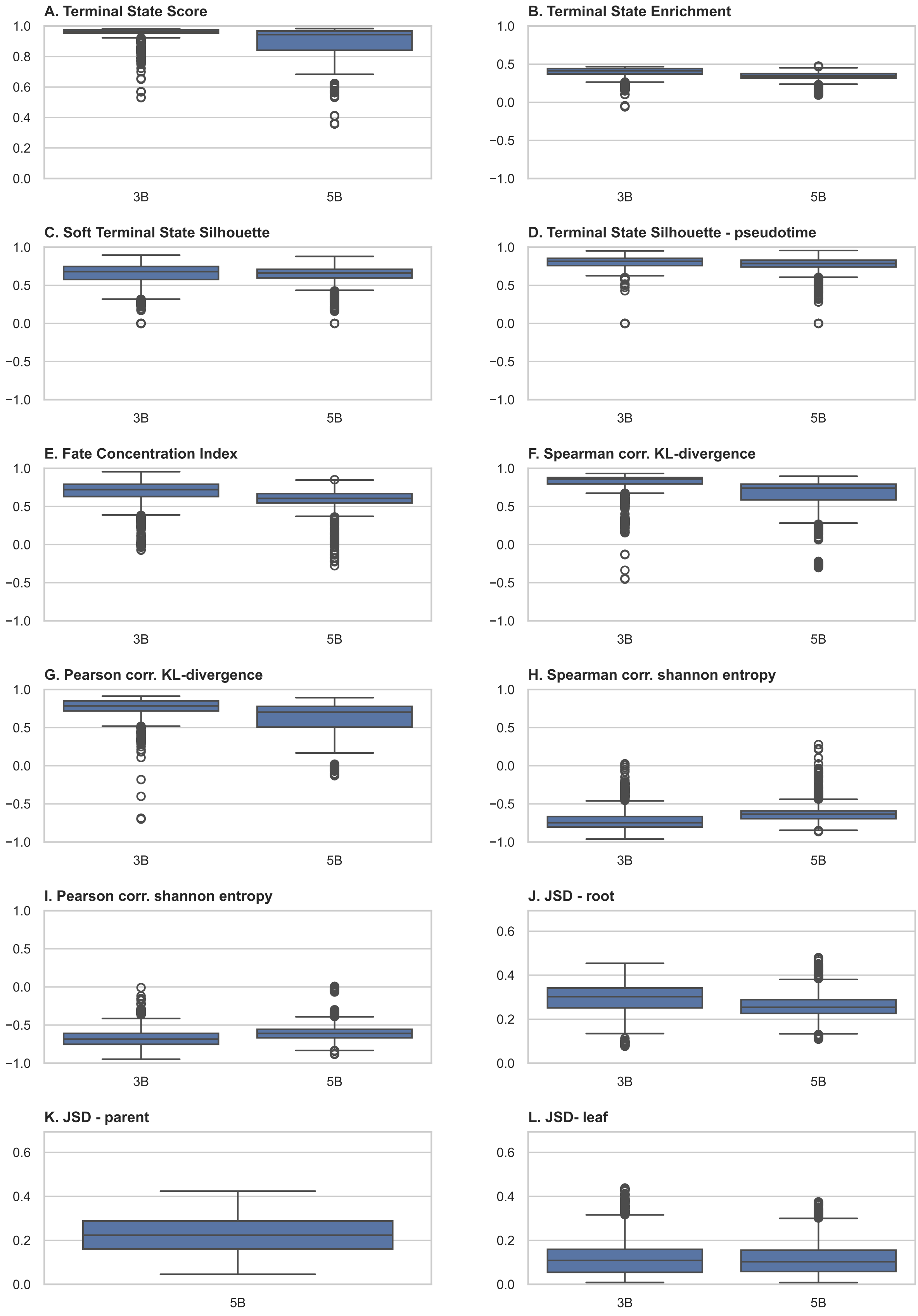


**Supplementary Figure 4.** Evaluation of ATLAS performance with Cellrank as TI strategy on the synthetic data.

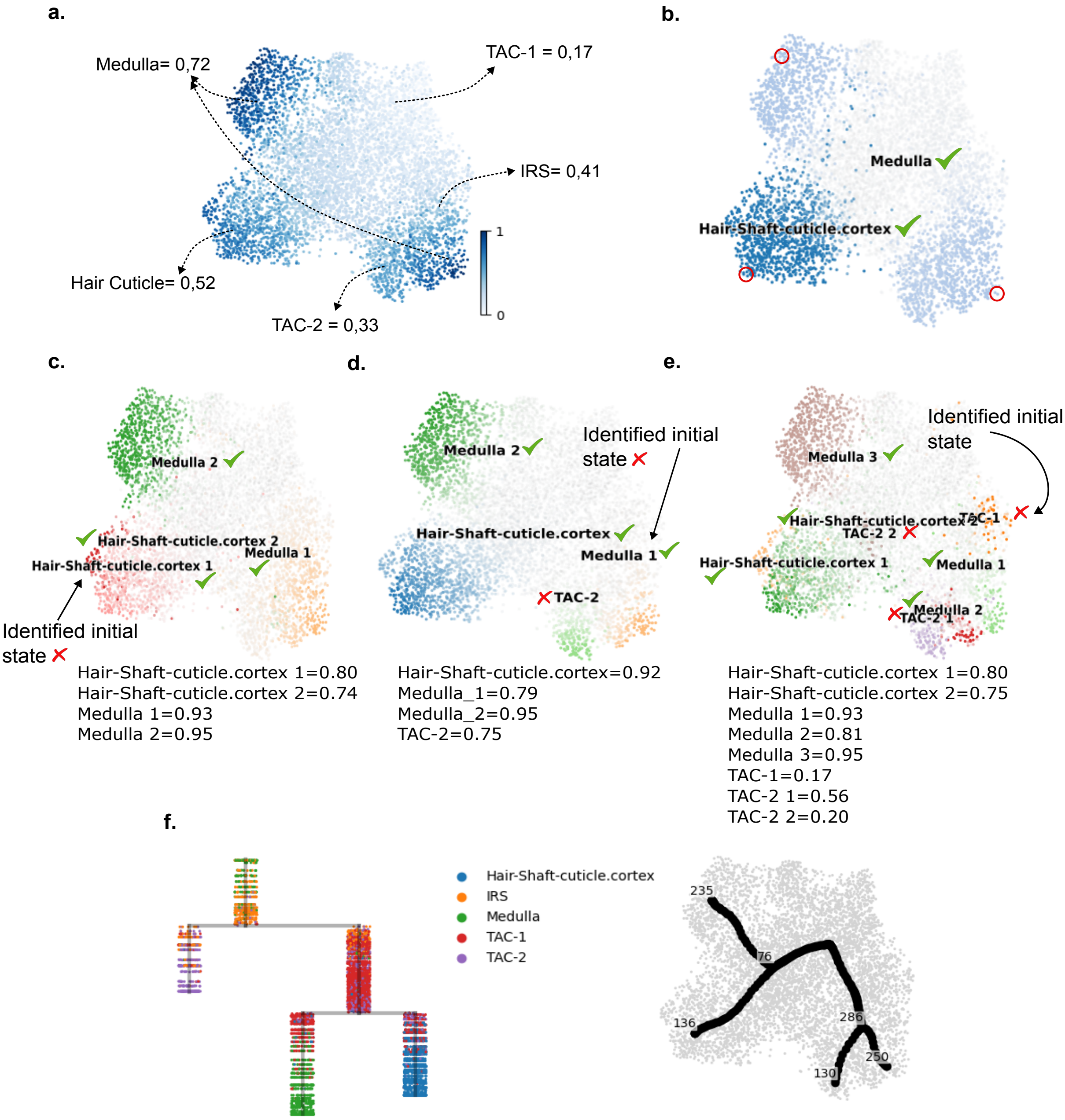


**Supplementary Figure 5.** Supplementary images for the SHARE-seq mouse hair dataset. **(a)** Pseudotime inferred from scRNA-seq data only and average pseudotime per celltype . **(b)** Inferred developmental trajectories using the original Palantir method on scRNA-seq data only. **(c)** Inferred developmental trajectories using the original CellRank `PseudotimeKernel` method on scRNA-seq data only, with $n_{\text{macrostates}} = 4$. The initial inferred state is highlighted. **(d)** Inferred developmental trajectories using ATLAS with $n_{\text{macrostates}} = 4$. The initial inferred state is highlighted. **(e)** Inferred developmental trajectories using the original CellRank `PseudotimeKernel` method on scRNA-seq data only, with $n_{\text{macrostates}} = 8$. The initial inferred state is highlighted. **(f)** Dendrogram and developmental graph obtained using ATLAS with $n_{\text{macrostates}} = 4$. Cells are colored according to their cluster annotation.

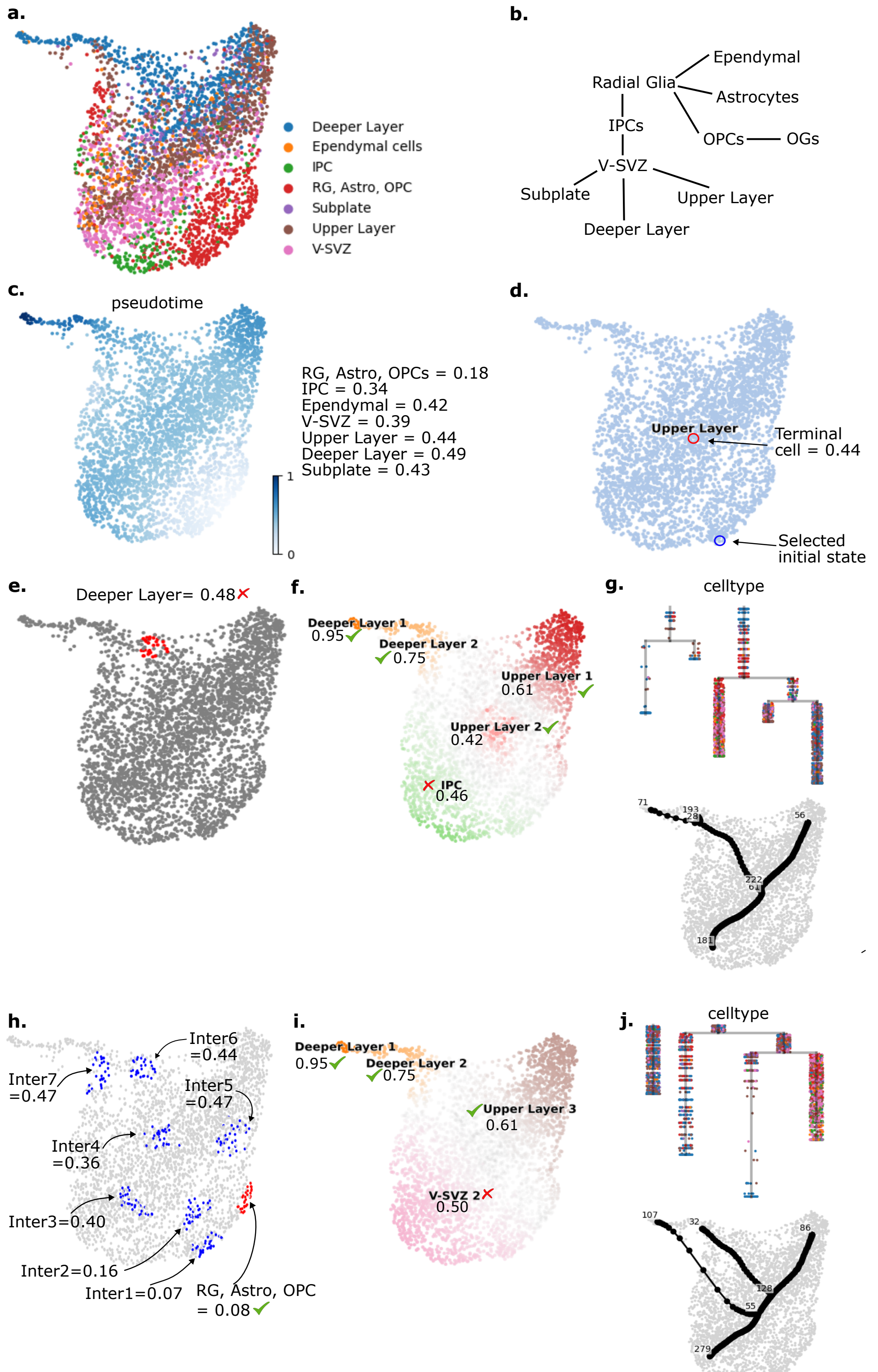


**Supplementary Figure 6.** Results of ATLAS on the Embryonic E18 Mouse Brain dataset. **(a)** Cell type annotations. **(b)** Expected developmental tree. **(c)** Inferred pseudotime and average pseudotime per cell cluster. **(d)** Inferred terminal state and developmental lineages using ATLAS and Palantir as undelying TI framework. **(e)** Inferred initial macrostate using CellRank as underlying TI framework with $n_{\text{macrostates}} = 6$. Average macrostate pseudotime is displayed. **(f)** Inferred terminal macrostates and developmental lineages using CellRank as underlying TI framework with $n_{\text{macrostates}} = 6$. Average terminal macrostate pseudotime is displayed. **(g)** Developmental graph and dendrogram obtained using scFates on the fate probabilities in plot **f**. Cells are colored according to their cluster. **(h)** Inferred initial and intermediate macrostates using CellRank as underlying TI framework with $n_{\text{macrostates}} = 12$. Average macrostate pseudotime is displayed. **(i)** Inferred terminal macrostates and developmental lineages using CellRank as underlying TI framework with $n_{\text{macrostates}} = 12$. Average terminal macrostate pseudotime is displayed. **(j)** Developmental graph and dendrogram obtained using scFates on the fate probabilities in plot **i**. Cells are colored according to their cluster.

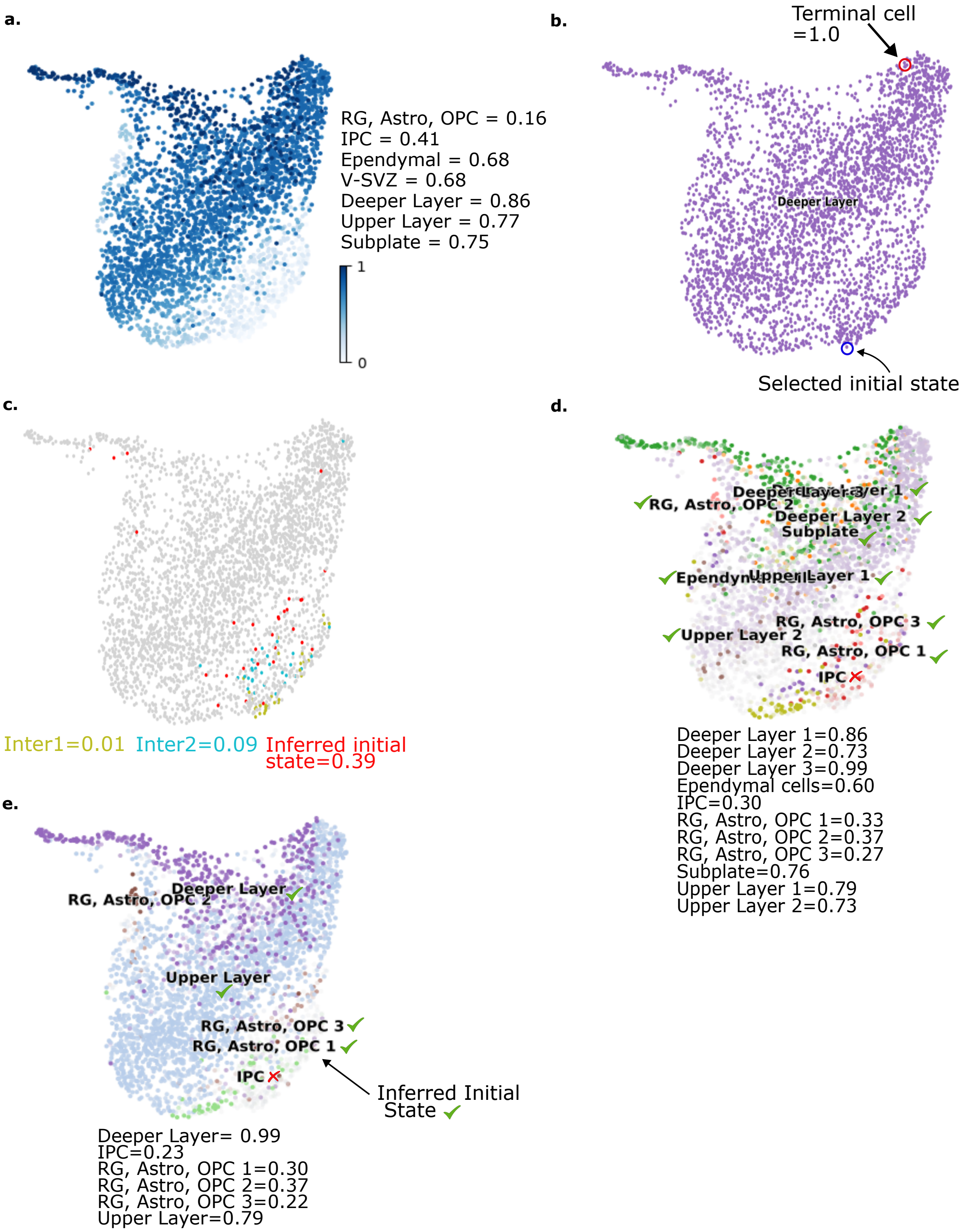


**Supplementary Figure 7.** Results of trajectory inference algorithms on the Embryonic E18 Mouse Brain dataset using scRNA-seq only data. **(a)** Inferred pseudotime and average pseudotime per cell cluster. **(b)** Inferred terminal state and developmental lineages using the original Palantir algorithm. **(c)** Inferred initial and intermediate macrostates using CellRank `PseudotimeKernel` when $n_{\text{macrostates}} = 12$. Average macrostate pseudotime is displayed. **(d)** Inferred developmental lineages and terminal macrostates using CellRank `PseudotimeKernel` when $n_{\text{macrostates}} = 12$. Average pseudotime per terminal macrostate is displayed. **(e)** Inferred developmental lineages, initial and terminal macrostates using CellRank `PseudotimeKernel` when $n_{\text{macrostates}} = 6$. Average pseudotime per terminal macrostate is displayed.

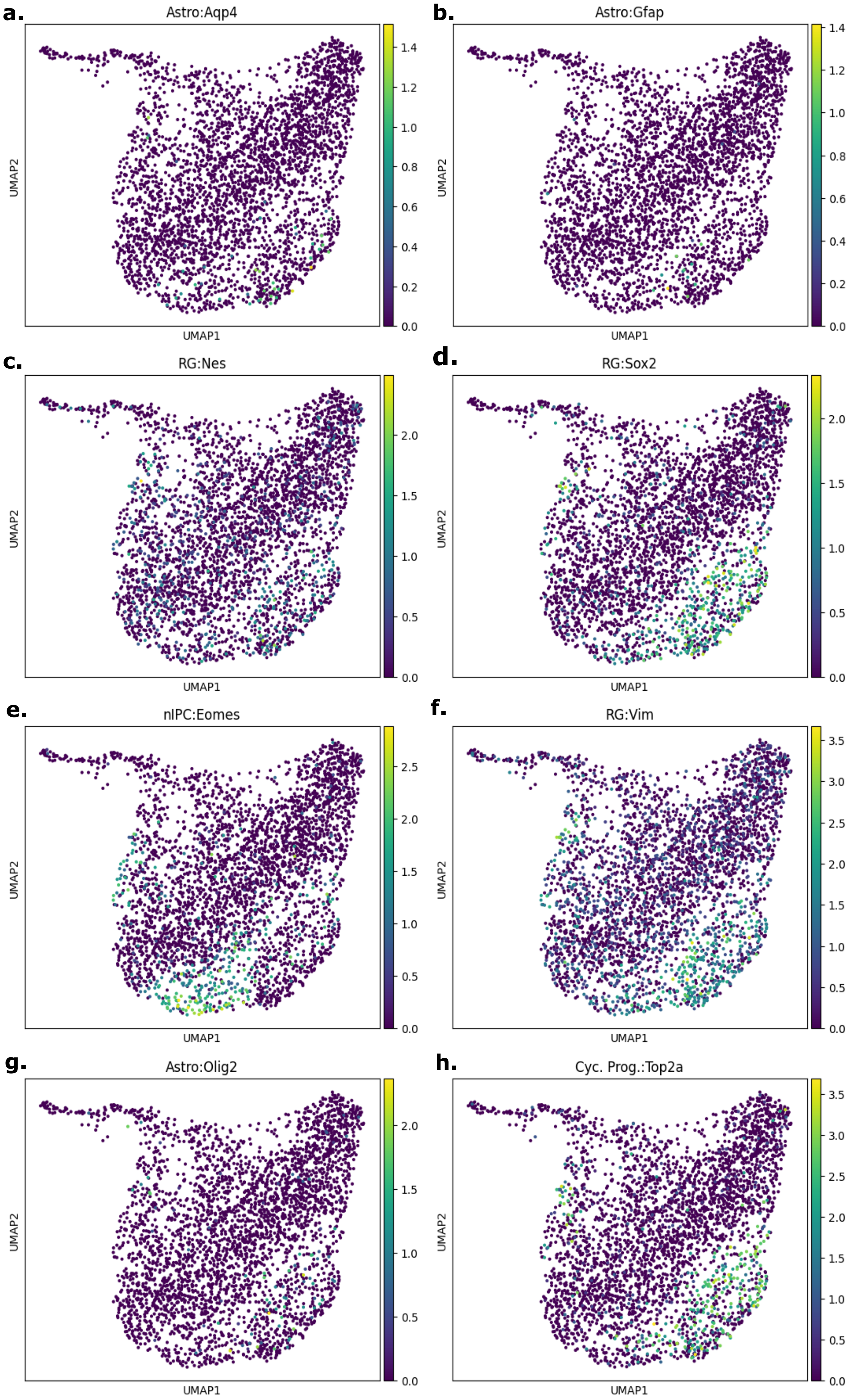


**Supplementary Figure 8.** Fresh Embryonic E18 Mouse Brain Dataset- expression of known marker genes.

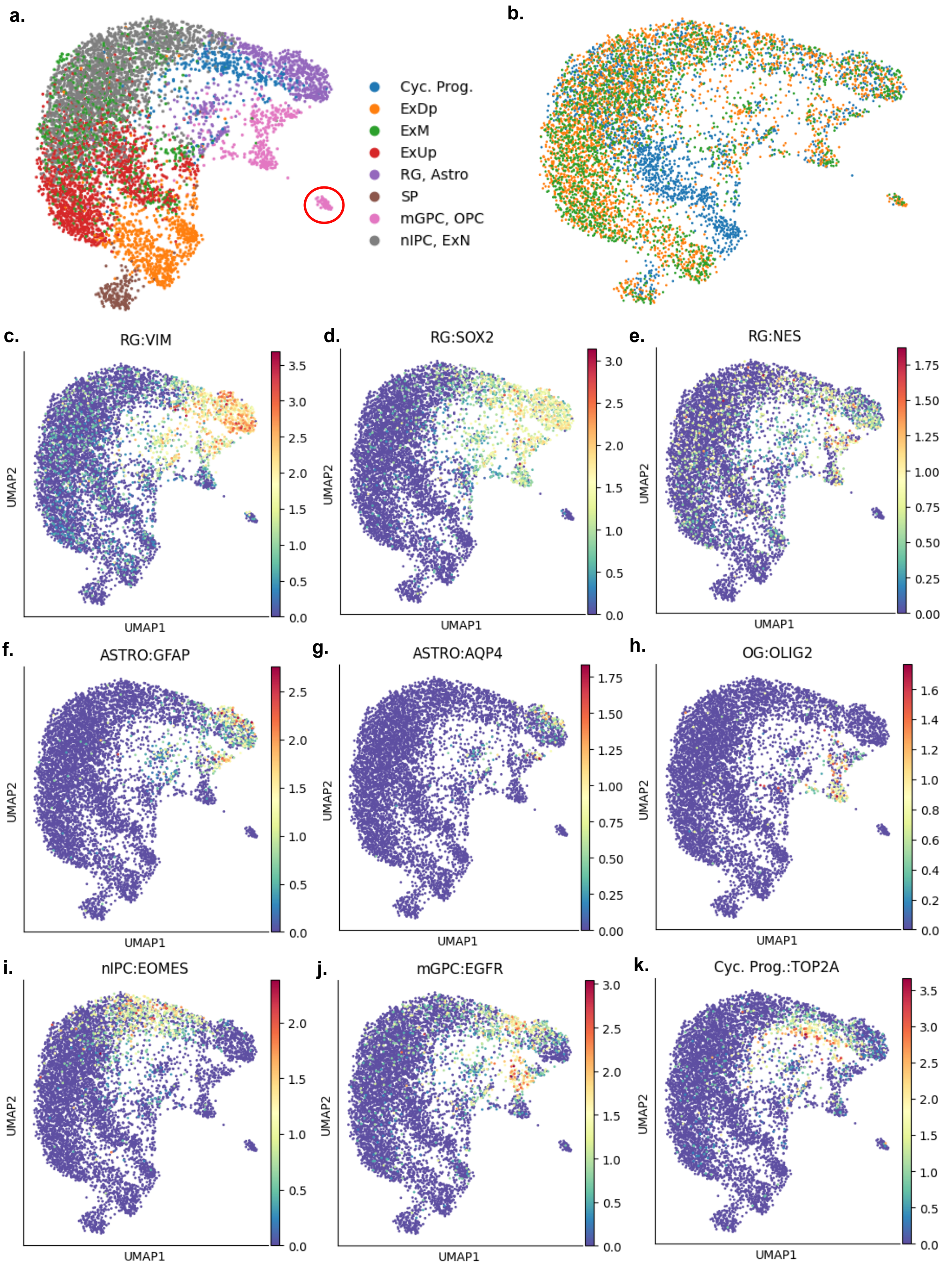


**Supplementary Figure 9.** Human Fetal Brain dataset. **(a)** Cell type annotation. The red circle highlights disconnected cells. **(b)** Sample annotation. **(c-k)** Expression of known marker genes.

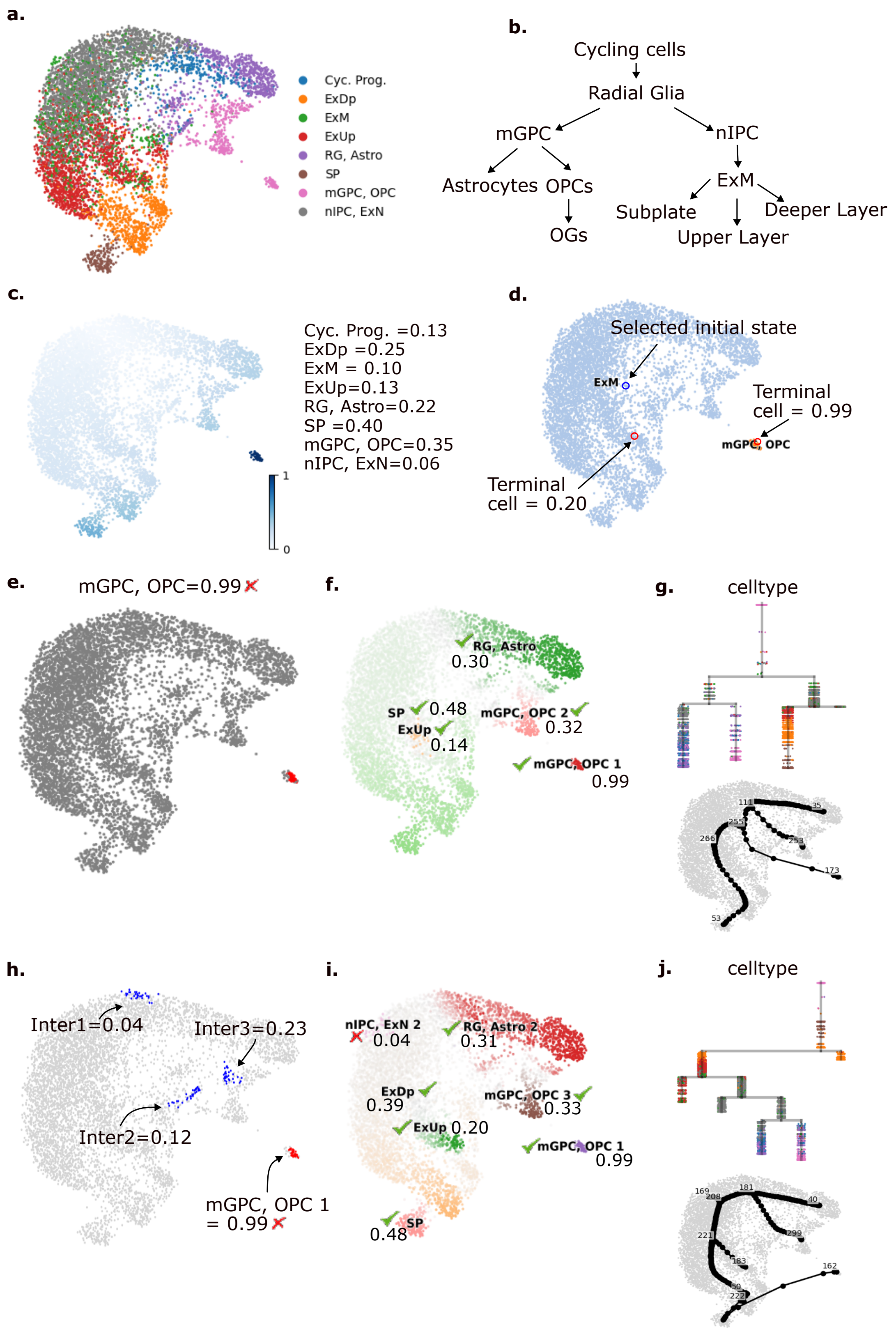


**Supplementary Figure 10.** Results of ATLAS on the Human Fetal Brain dataset. **(a)** Cell type annotations. **(b)** Expected developmental tree. **(c)** Inferred pseudotime and average pseudotime per cell cluster. **(d)** Inferred terminal states and developmental lineages using ATLAS and Palantir as underlying TI framework. **(e)** Inferred initial macrostate and average pseudotime using CellRank as underlying TI framework with $n_{\text{macrostates}} = 5$. **(f)** Inferred terminal macrostates, average pseudotime and developmental lineages using CellRank as underlying TI framework with $n_{\text{macrostates}} = 5$. **(g)** Developmental graph and dendrogram obtained using scFates on the fate probabilities from plot **f**. Cells are colored according to their cluster. **(h)** Inferred initial and intermediate macrostates with avrage pseudotime using CellRank as underlying TI framework with $n_{\text{macrostates}} = 10$. **(i)** Inferred terminal macrostates, average pseudotime and developmental lineages using CellRank as underlying TI framework with $n_{\text{macrostates}} = 10$. **(j)** Developmental graph and dendrogram obtained using scFates on the fate probabilities in plot **i**. Cells are colored according to their cluster.

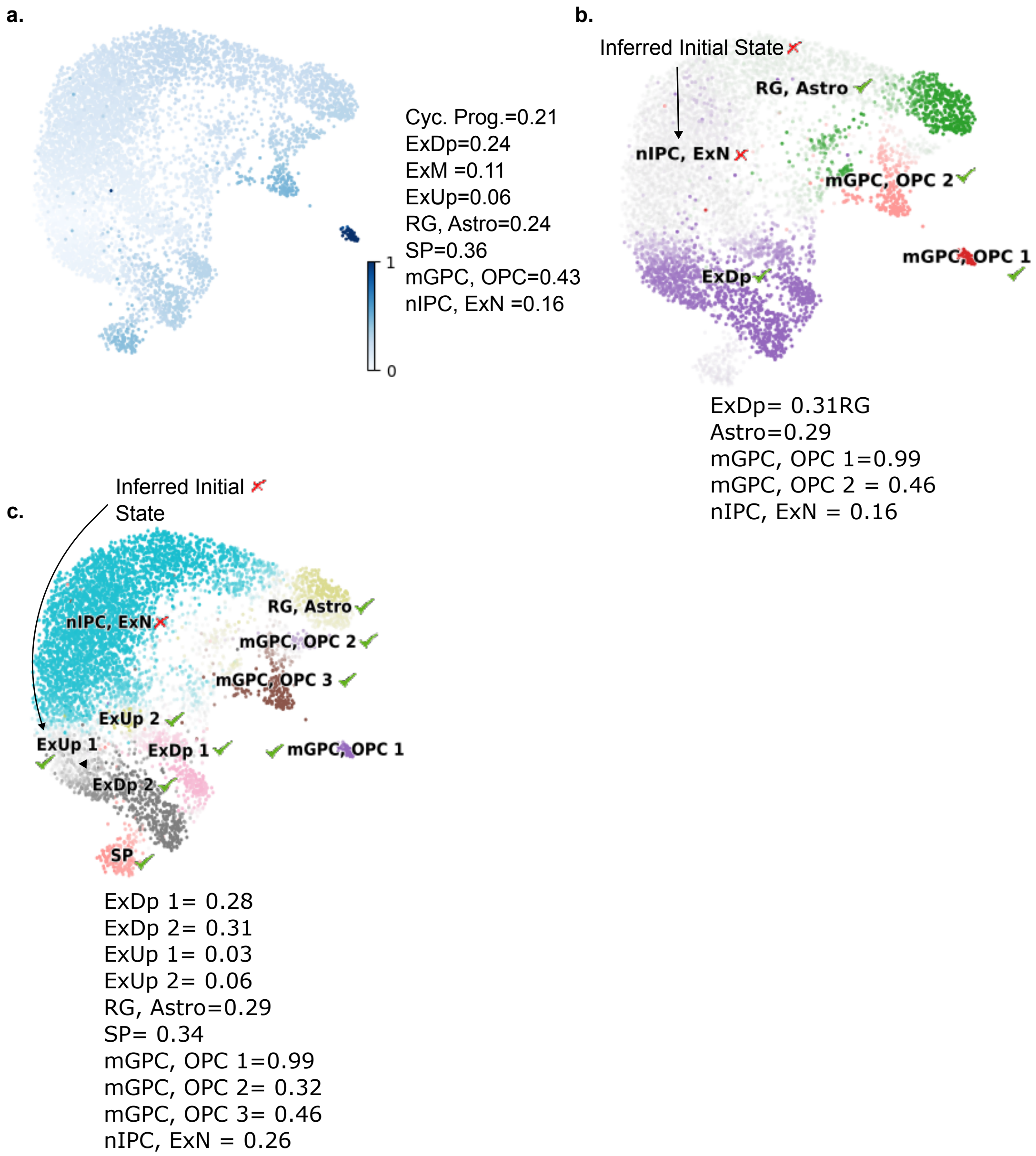


**Supplementary Figure 11.** Results of trajectory inference algorithms on the Human Fetal Brain dataset using scRNA-seq only data. **(a)** Inferred pseudotime and average pseudotime per cell cluster. **(b)** Inferred initial state, developmental lineages and terminal macrostates using CellRank `PseudotimeKernel` when $n_{\mathrm{macrostates}} = 5$. Average pseudotime per terminal macrostate is displayed. **(c)** Inferred developmental lineages, initial and terminal macrostates using CellRank `PseudotimeKernel` when $n_{\mathrm{macrostates}} = 10$. Average pseudotime per terminal macrostate is displayed.

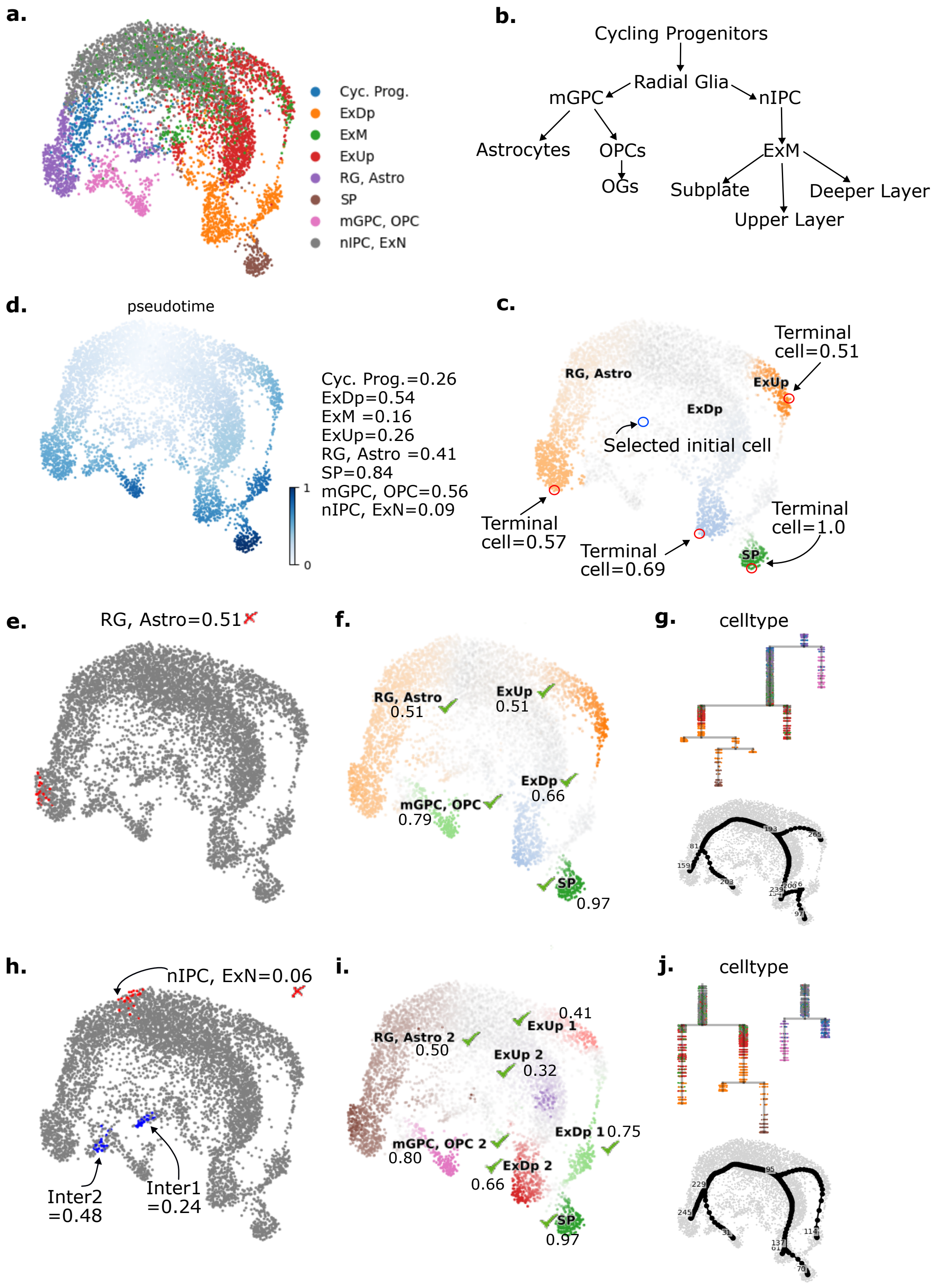


**Supplementary Figure 12.** Results of ATLAS on the Human Fetal Brain dataset, without disconnected cells. **(a)** Cell type annotations. **(b)** Expected developmental tree. **(c)** Inferred pseudotime and average pseudotime per cell cluster. **(d)** Inferred terminal states and developmental lineages using ATLAS and Palantir as underlying TI framework. **(e)** Inferred initial macrostate and average pseudotime using CellRank as underlying TI framework with $n_{\text{macrostates}} = 5$. **(f)** Inferred terminal macrostates, average pseudotime and developmental lineages using CellRank as underlying TI framework with $n_{\text{macrostates}} = 5$. **(g)** Developmental graph and dendrogram obtained using scFates on the fate probabilities from plot **f** Cells are colored accordind to their cluster. **(h)** Inferred initial and intermediate macrostates with average pseudotime using CellRank as underlying TI framework with $n_{\text{macrostates}} = 10$. **(i)** Inferred terminal macrostates, average pseudotime and developmental lineages using CellRank as underlying TI framework with $n_{\text{macrostates}} = 10$. **(j)** Developmental graph and dendrogram obtained using scFates on the fate probabilities in plot **i**. Cells are colored according to their cluster.

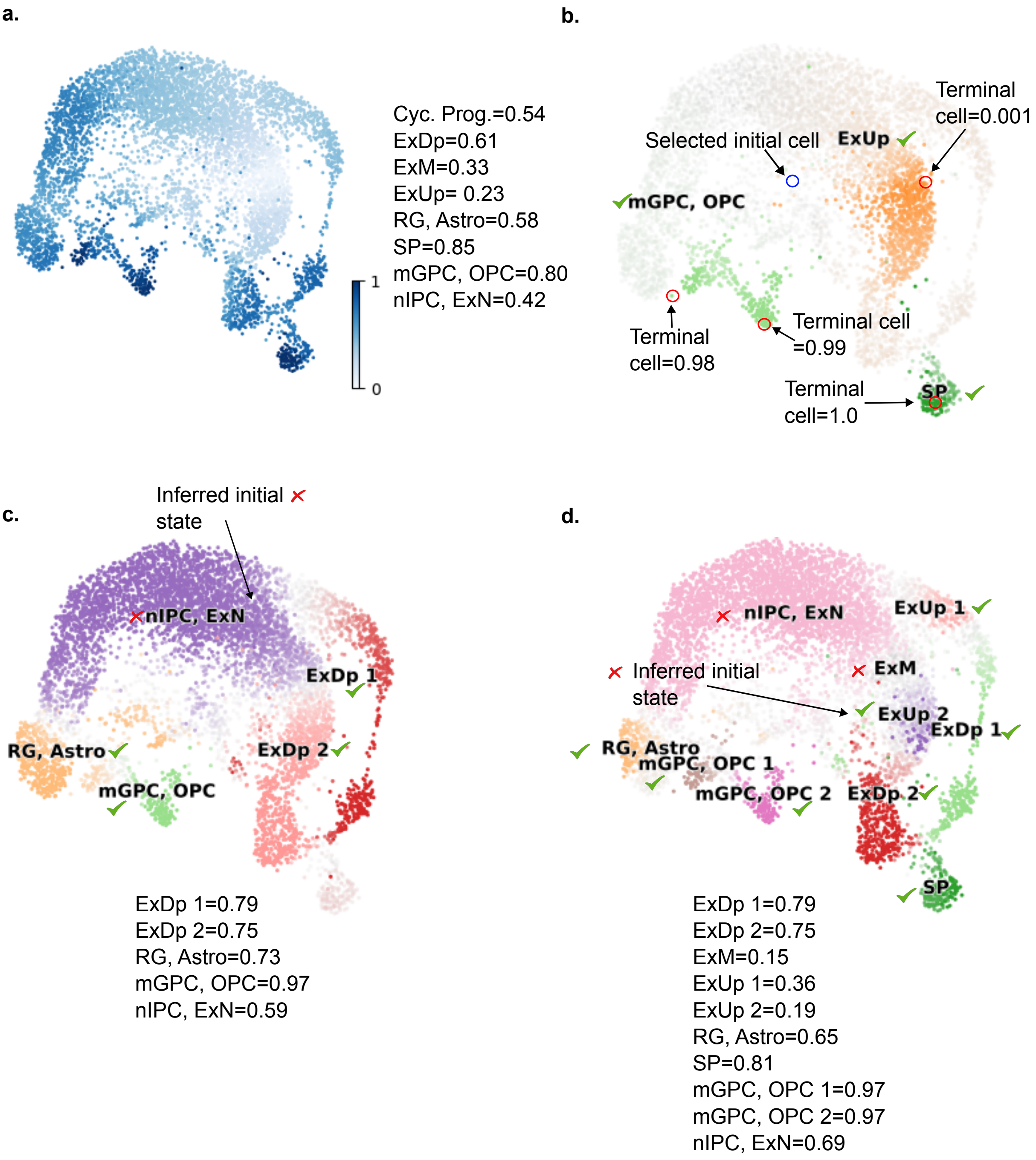


**Supplementary Figure 13.** Results of trajectory inference algorithms on the Fetal Human Brain dataset using scRNA-seq only data, without disconnected cells. **(a)** Inferred pseudotime and average pseudotime per cell cluster. **(b)** Inferred terminal cells and developmental lineages using Palantir. Pseudotime per terminal cell is displayed. **(c)** Inferred initial state, developmental lineages and terminal macrostates using CellRank `PseudotimeKernel` when $n_{\text{macrostates}} = 5$. Average pseudotime per terminal macrostate is displayed. **(d)** Inferred developmental lineages, initial and terminal macrostates using CellRank `PseudotimeKernel` when $n_{\text{macrostates}} = 10$. Average pseudotime per terminal macrostate is displayed.
